# Self-organized Recovery of Coordinated Locomotion in Crickets via Prosthetic Limb Integration

**DOI:** 10.64898/2026.01.15.699627

**Authors:** Dai Owaki, Hitoshi Aonuma

## Abstract

Distributed sensorimotor interactions facilitate the coordination of multi-legged locomotion in insects without centralized control, yet the mechanisms that allow coordinated locomotion to re-emerge following limb loss remain poorly understood. Here, we systematically evaluate the effects of leg amputation and the integration of prosthetic legs on walking coordination in crickets *Gryllus bimaculatus*. Spherical treadmill experiments revealed that leg amputation disrupts inter-leg phase coupling, decreases locomotor speed, and alters spatial foot placement in a state-dependent manner, indicating impaired load-mediated coordination. Prosthetic legs did not merely restore intact kinematics; instead, they selectively reinstated coherent temporal coordination and axis-specific spatial organization. This structured recovery illustrates that re-introducing mechanically relevant sensory constraints is sufficient to re-engage distributed coordination networks, even in the absence of anatomical integrity. The selective recovery of temporal over spatial coordination, demonstrated here quantitatively for the first time, reveals the hierarchical architecture of distributed locomotion control and elucidates an embodied principle by which sensory–mechanical feedback facilitates the self-organization of resilient multi-legged locomotion following morphological intervention. Load-mediated sensory signals thus emerge as a key driver of distributed coordination in biohybrid systems, providing design principles for adaptive prosthetic engineering and sensorimotor rehabilitation.

## 1 Introduction

Understanding animal locomotion is a fundamental scientific challenge that intersects biology, neuroscience, and robotics. Insects, in particular, serve as powerful model systems; despite their relatively simple nervous systems [1–6], they have evolved remarkably flexible and adaptive walking behaviors across diverse environments. Central to these capabilities are the principles of *embodied intelligence* [7, 8], where neural circuits, body morphology, and environmental sensory feedback interact in tightly integrated way [9]. Classical studies of insect locomotion have elucidated the regulatory mechanisms underlying inter-leg coordination [10–12], demonstrating how multi-legged animals achieve both stability and agility through distributed control strategies [13, 14]. Beyond advancing neuroethology, insights derived from insect locomotion have significantly influenced biorobotics [15, 16] and the formulation of general principles of locomotor control [17, 18].

A striking example of embodied robustness is the capacity of insects to maintain locomotion even after losing one or more legs. Foundational studies in cockroaches *Blatta orientalis* [19] and stick insects *Carausius morosus* [20, 21] revealed that leg amputation alters gait patterns; however, inter-leg coordination adapts flexibly to preserve walking performance. Similarly, desert ants *Cataglyphis* exhibit remarkable morphological adaptability: when their leg length is experimentally altered using “stilts” or “stumps,” they adjust their walking pattern accordingly, illustrating that even changes in body morphology can be accommodated by flexible locomotor control [22, 23]. More recent research in fruit flies *Drosophila* [24] and crickets *Gryllus bimaculatus* [25] has demonstrated that neuromuscular systems sustain coordinated locomotion following amputation by modulating speed and muscle activity in the remaining legs. These findings underscore that insect locomotion is not solely dictated by neural circuitry; rather, it emerges from the interplay of morphology and sensory feedback, endowing the system with exceptional resilience [9].

Recent advances in biohybrid methodologies have opened new avenues for restoring and modulating locomotion through the integration of artificial materials with living tissues. For instance, the leg muscles of beetles *Mecynorrhina torquata* can be electrically stimulated to regulate flight and walking behaviors [26–28]; similar bioelectrical actuation has also been applied to the ring muscles of jellyfish *Aurelia aurita/coerulea* to facilitate swimming locomotion [29–31]. In a complementary line of work, insect–machine hybrid systems have been developed to interface the neural circuits of the silkmoth *Bombyx mori* with robotic platforms, allowing quantitative investigation of sensory–motor adaptability and neural control under semi-natural conditions [32, 33]. Three-dimensional printed prosthetic limbs attached to Madagascar cockroaches *Gromphadorhina Portentosa* have enhanced self-righting and load-bearing behaviors [34]. Furthermore, the development of *Xenobots*, bioengineered constructs that exploit the self-healing and motile properties of *Xenopus laevis* cells, demonstrates how biological substrates can be programmed to perform goal-directed movements [35, 36]. These accomplishments highlight a growing convergence between biomechanics, neuroscience, and robotics, pointing toward a unified framework in which motor control and recovery can be understood as emergent properties of embodied systems.

A key unresolved question is whether prosthetic-mediated locomotor recovery occurs uniformly across the temporal and spatial dimensions of gait or whether these dimensions dissociate because of the distinct functional properties of their underlying mechanosensory pathways. The most relevant precedent is the study by Noah et al. (2004), which showed that cockroaches equipped with rigid “peg legs” (passive replacements lacking distal sensory input) could recover coordinated locomotion. This finding implicated load-mediated ground contact as a minimal mechanical cue sufficient to re-engage distributed inter-leg coordination [37]. However, that study did not determine whether temporal inter-leg phase coupling and spatial foot-placement accuracy recovered to comparable extents, nor did it identify the mechanosensory pathways underlying each dimension of recovery. This distinction is mechanistically important: temporal coordination primarily depends on load-sensitive mechanoreceptors and thoracic central pattern generators, whereas spatial accuracy additionally requires intact joint-angle proprioception, which cannot be restored by a passive prosthetic limb. Resolving this question requires an experimental preparation in which inter-ganglion coordination across multiple leg combinations can be quantified independently. The cricket, *Gryllus bimaculatus*, is particularly well suited to this purpose: its six legs are governed by three thoracic ganglia, each of which controls a bilaterally symmetric pair of legs, allowing temporal and spatial coordination across fore-, mid-, and hindlimb combinations to be quantified independently and simultaneously within a single experimental subject.

Resolving this dissociation also addresses a central engineering challenge in biohybrid prosthetics and sensorimotor rehabilitation. A mechanically passive prosthesis that restores only load-mediated ground-contact signals establishes a lower bound for prosthetic efficacy: the coordination that recovers under these minimal constraints reveals what can be achieved without active sensory replication. Conversely, the gait dimensions that remain impaired identify the specific sensory functions, such as joint-angle proprioception, that must be incorporated into next-generation biohybrid prosthetic devices to achieve more complete locomotor rehabilitation. The cricket prosthetics paradigm therefore serves not merely as a neurobiological preparation but also as a living proof of concept for hierarchical recovery strategies that may inform the design of adaptive prosthetic limbs and sensorimotor rehabilitation protocols in biohybrid systems.

The present study addresses these open questions by demonstrating, quantitatively for the first time, a spatiotemporal dissociation in prosthetic locomotor recovery. Temporal inter-leg phase coordination recovers selectively and substantially following prosthetic integration, whereas spatial foot-placement trajectories recover only partially. This dissociation is not merely phenomenological. It is predicted *a priori* by the functional asymmetry between campaniform sensilla-mediated load-timing signals, which are preserved because the prosthetic leg contacts the substrate and reinstates load-onset and load-release events, and chordotonal organ-mediated joint-angle proprioception, which is absent because the prosthesis lacks articulated joints. The five-condition within-subject design (Intact → Amp,1 → Proth,1 → Amp,2 → Proth,2) and DeepLabCut-based multi-dimensional kinematic analysis together provide the experimental resolution necessary to establish this spatiotemporal dissociation for the first time.

Building on this perspective, our objective was to translate the adaptive principles of natural insect locomotion into a functional biohybrid system aimed at restoring locomotion. To achieve this, we investigated the impact of leg amputation on the spatiotemporal gait coordination in crickets (*Gryllus bimaculatus*) and evaluated whether integrating prosthetic legs into their body morphology could re-establish coordinated walking. Utilizing a spherical treadmill, we systematically analyzed the locomotor patterns of intact, amputated, and prosthesis-attached crickets, quantifying spatiotemporal modifications in inter-leg coordination through a deep-learning-based markerless tracking algorithm (DeepLabCut: DLC) [38, 39]. Collectively, these experiments helped us understand how the loss and subsequent re-introduction of mechanically relevant sensory–motor pathways influence locomotor coordination across multiple dynamical scales. By directly comparing intact, amputated, and prosthesis-integrated conditions, we demonstrate that body-integrated prosthetic legs selectively restore coherent temporal and spatial characteristics of insect walking, rather than merely reinstating intact kinematics. This research establishes a biohybrid framework for investigating and reconstructing embodied locomotor control through targeted morphological interventions.

## 2 Materials and Methods

### 2.1 Intervention Experiment Utilizing Leg Amputation and Prosthetic Leg Implementation

We investigated how leg amputation disrupts spatiotemporal coordination during walking in the cricket *Gryllus bimaculatus* (Fig. 1 A) [25] and whether a body-integrated prosthetic leg (Fig. 1 B) can facilitate the restoration of coordinated locomotion. We employed a spherical treadmill (Fig. 1 C and Movie SM1) to conduct a comprehensive within-framework analysis of locomotor patterns across five distinct experimental conditions (Fig. 1 D): (a) intact animals; (b) right middle leg amputation at the femur-tibia (FTi) joint; (c) prosthetic attachment at the right FTi joint; (d) bilateral amputation of both left and right middle legs at the FTi joints; and (e) prosthetic legs affixed to both FTi joints. This experimental framework allowed us to differentiate the consequences of leg amputation from those associated with morphological reintegration, thereby facilitating a direct assessment of the prosthetic leg’s effectiveness in re-establishing coordinated walking patterns. The present study specifically investigates walking coordination recovery following amputation and prosthetic replacement of the middle legs. The effects of equivalent interventions at the foreleg or hindleg positions are expected to differ substantially because each leg pair serves a distinct functional role in hexapod locomotion. The forelegs primarily support steering and substrate exploration, the middle legs serve as the primary load-bearing structures and hubs for inter-ganglionic coordination, and the hindlegs primarily generate propulsive force [12, 14]. Systematic investigation across all leg positions therefore represents an important direction for future work. We quantified changes in inter-leg coordination using a deep-learning–based markerless tracking methodology (DeepLabCut, DLC) [38, 39], which enables high-resolution and unbiased evaluation of gait dynamics. Representative walking patterns for each experimental condition are shown in the corresponding videos (Movies SM2–SM11) and are quantitatively characterized and visualized in Figs. S5–S9.

**Figure 1:**
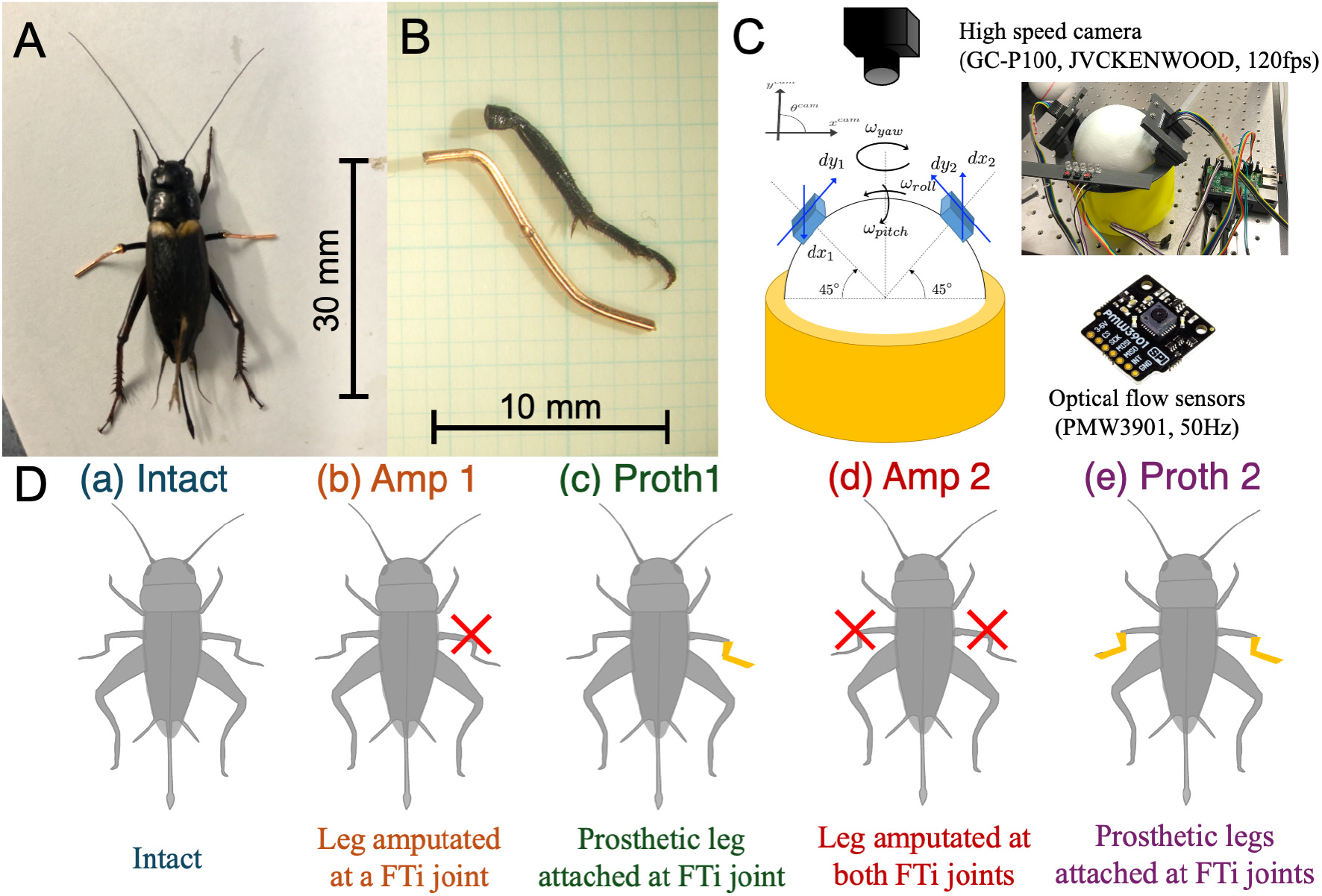
Overview of Experimental Manipulations Using Leg Amputation and Prosthetic Leg Implementation: (**A**) *Gryllus bimaculatus* (female). (**B**) Artificial prosthetic leg (2.76 ± 0.18 mg) that replicates a cricket middle leg, using a deformable metal wire. (**C**) Measurement system (Movie SM1) of walking velocity and angular velocity of body rotation by using two optical flow sensors. (**D**) Experimental conditions (a) Intact; (b) Leg amputated at the right FTi joint; (c) Prosthetic leg attached at the right FTi joint; (d) Leg amputated at both FTi joints; and (e) Prosthetic legs attached at both FTi joints. Representative walking patterns for each experimental condition are shown in the corresponding videos (Movies SM2–SM11) and are quantitatively characterized and visualized in Figs. S5–S9.

### 2.2 Animals

The crickets *Gryllus bimaculatus* were raised in a laboratory colony under a 14h:10h light and dark cycle at 28 ± 2 ^◦^C (with lights on at 6:00). They were fed insect food pellets (Oriental Yeast Co., Tokyo, Japan) and had access to water ad libitum. Only adult female crickets that had molted within the previous two weeks were used in this study.

### 2.3 Ethics

Research involving insects (invertebrates) at Tohoku University and Kobe University is not currently governed by institutional animal care regulations. However, we fully endorse the ethical considerations and responsibilities inherent in animal research, regardless of whether formal approval is required for invertebrates or mandated for vertebrates [40–42]. In our cricket experiments, we adhered to the principles of harm minimization and precaution, as well as the 4Rs framework (reduction, replacement, refinement, and reproduction). Furthermore, we recognize that future research on biohybrid systems, particularly those that may challenge ethical boundaries not yet comprehensively addressed by ethicists or legislators, requires careful evaluation of both animal welfare and broader societal implications [41, 42].

### 2.4 Behavioral Experiments

Randomly selected crickets (*N* = 5) from the colony were used in the experiments. Before being placed on a treadmill to observe their walking gait patterns, each cricket was anesthetized using CO^2^ gas. The treadmill, made from a Styrofoam sphere (100 mm in diameter) supported by an upward air stream, facilitated the observation (Fig.1C). A 2 mm diameter plastic rod, created using a 3D printer (Method X Carbon Fiber Edition; Carbon, Redwood City, CA, USA), was attached to the thorax of each cricket with a UV-curable liquid plastic adhesive (Bondic refill cartridge; Bondic, Aurora, CO, USA). This rod was connected to a ring mounted on a manipulator, allowing for precise positioning of the cricket on the Styrofoam sphere. This setup enabled the cricket to walk freely while allowing adjustments in its orientation and ground clearance (Movie SM1). This tethered spherical treadmill setup is an established standard for high-resolution locomotion recording in insects [43] and enabled simultaneous quantification of translational and rotational velocity components alongside individual limb kinematics within the same within-subject design.

Kinematic recordings were collected from five adult female crickets (cricket017–021 in Table S3). The experimental conditions (a)–(e), illustrated in Fig. 1D, were conducted sequentially on a single subject following this protocol. (a) Intact crickets, without any interventions, were placed on the treadmill, and their walking patterns were recorded as controls; they were subsequently removed from the treadmill for surgical intervention. (b) The cricket was re-anesthetized with CO^2^ gas, and the right middle leg was amputated at the femur-tibia (FTi) joint using fine scissors. The cricket was then returned to the treadmill to record its walking gait. (c) After removal from the treadmill, the cricket was anesthetized again, and a prosthetic leg (Fig. 1B) was inserted into the cuticle at the amputated FTi joint, secured with UV-curable liquid to restore appropriate leg morphology. (d) Following anesthesia, the prosthetic right middle leg applied in condition (c) was removed, the left middle leg was also amputated at the FTi joint as in (b), and gait patterns were recorded during bilateral middle-leg amputation on the treadmill. (e) Similarly to (c), prosthetic legs were inserted into the cuticles of both amputated middle-leg FTi joints (Fig. 1A), fixed with UV-curable liquid, and gait patterns were recorded on the treadmill. To control for potential effects of mass addition, a subset of animals underwent an additional condition in which a mass-matched inert attachment lacking prosthetic leg geometry was fixed to the amputation stump; representative walking sequences are provided in Supplementary Movie SM12 and SM13.

We quantified the spatiotemporal leg coordination patterns, locomotor velocity, and body rotational angular velocity under each condition. To achieve this, we utilized a high-speed camera (GC-P100, JVCKENWOOD Corporation; 120 fps) and two-dimensional optical-flow sensors (PMW3901, Pimoroni; 50 Hz) (Fig. 1C). Motion data of the crickets (videos) were stored on an SD memory card within the camera. The rotational angular velocities of the spherical treadmill along the yaw, roll, and pitch axes were measured using two optical-flow sensors mounted on the treadmill base (Fig. 1C). Sensor signals were quantified and recorded via SPI (Serial Peripheral Interface) communication using a Raspberry Pi 4B+ (Raspberry Pi Foundation). To synchronize video recordings with optical-flow measurements, small light-emitting diodes (LEDs) served as temporal markers. By triggering LED blinking with tactile switches and logging the corresponding time-series data on the Raspberry Pi, we achieved precise synchronization between motion images and sensor signals for subsequent gait analysis (Fig. S3).

### 2.5 Gait Analysis

To quantify leg coordination patterns from two-dimensional high-speed camera recordings (640 × 320 pixels at 120 fps), we employed a deep learning-based pose estimation algorithm, DeepLabCut (DLC) [38, 39]. By annotating training datasets comprising 50–200 frames per condition, the algorithm automatically tracked 21 anatomical and experimental landmarks (Head, Pro, Meso, Meta, LF1, LF2, LM1, LM2, LH1, LH2, RF1, RF2, RM1, RM2, RH1, RH2, Bar, Axis, Fix, LED1, and LED2) during each trial [25] (Figs. S2, S3 and Movies SM2-SM11). The pose estimation error after DLC processing was maintained below 2.0 pixels (out of 640 × 320 pixels), indicating sufficient accuracy for subsequent analyses.

Using the estimated positions, we quantified leg kinematics during walking to characterize inter-leg phase relationships, spatial foot trajectories, and the timing of touchdown (TD) and liftoff (LO) events (Fig. S4). DLC-derived marker trajectories were converted into time-series data for each leg endpoint (e.g., LF1 tarsus). From these trajectories, the instantaneous phase of each leg movement, *ϕ_i_*(*t*), was computed using the Hilbert transform [44], with the TD timing of each leg defined as the phase origin (*ϕ_i_* = 0). Inter-leg phase differences were quantified by calculating the relative phase (*ϕ_j_* − *ϕ_i_*) of each leg with respect to the TD timing of the left hind (LH) leg, enabling consistent comparisons of phase coupling across amputation and prosthetic conditions. Due to limitations inherent in two-dimensional kinematic recordings, TD and LO events were approximated using the anterior extreme position (AEP) and posterior extreme position (PEP) [11, 12] of each leg, respectively. For each leg (*i*), the vertical displacement time series of the tarsus, *y_i_*(*t*), was extracted. TD and LO events were detected independently for each leg using a peak-based method, where positive peaks in *y_i_*(*t*) were classified as TD events, while negative peaks (identified from −*y_i_*(*t*)) corresponded to LO events. Peak detection utilized a prominence threshold of 0.4 and a minimum inter-event interval of 24 frames (0.2 s), effectively separating stance–swing transitions across all experimental conditions. Sample sizes (*n*) for each condition correspond to the number of TD events of the LH leg, each representing an analyzed walking cycle. The resulting TD and LO indices were then used to extract the corresponding two-dimensional foot positions in normalized coordinates for further spatial analyses. Only bouts of sustained forward locomotion, identified by continuous forward displacement of the spherical treadmill, were included in the analysis; periods of immobility, erratic ball rotation, or apparent stumbling were excluded. No explicit inter-event-interval criterion was applied to exclude burst-stepping behavior that can occur in amputated insects [37, 45, 46].

Using signals from two two-dimensional optical flow sensors in Fig. 1C, we computed the rotational angular velocities of the spherical treadmill in yaw, roll, and pitch according to established methodologies [25, 43]. Based on the camera-frame coordinate definition, locomotor velocities in the (*x*) and (*y*) directions were calculated using the sphere radius (*l_s_* = 50 mm). Forward velocity was normalized by body length (BL) to account for inter-individual size variation, while angular velocity was retained with sign to preserve turning direction. To normalize (*x*)–(*y*) position and velocity data by cricket body size, all values were divided by the corresponding body length (BL; Table S3), resulting in unit conversions from mm or mm/s to BL or BL/s.

### 2.6 Statistical Analysis

To visualize phase-difference distributions, we constructed rose histograms using 36 equally spaced bins that spanned from 0 to 2*π* (Fig. 2). For each histogram, bin heights were normalized to represent probability densities, and the radial axes were scaled independently for each condition based on the maximum density observed across leg pairs. This approach allowed for accurate comparisons of distribution shapes while minimizing artifacts associated with fixed scaling. Mean resultant vectors were computed from the circular data to quantify the preferred phase direction and its concentration, following standard methods in circular statistics [47], and plotted as radial lines to indicate the preferred phase and its concentration. Differences in mean phase between the Intact condition and each manipulation were assessed using the Watson–Williams two-sample test, a circular analogue of ANOVA for comparing mean directions. For each comparison, we calculated the *F* -statistic, associated degrees of freedom, and corresponding *p*-value. Significance levels were reported as *p <* 0.01 (**), *p <* 0.05 (*), or non-significant (ns) (Table S1). Trials with fewer than two valid samples per condition were excluded from statistical evaluation.

**Figure 2:**
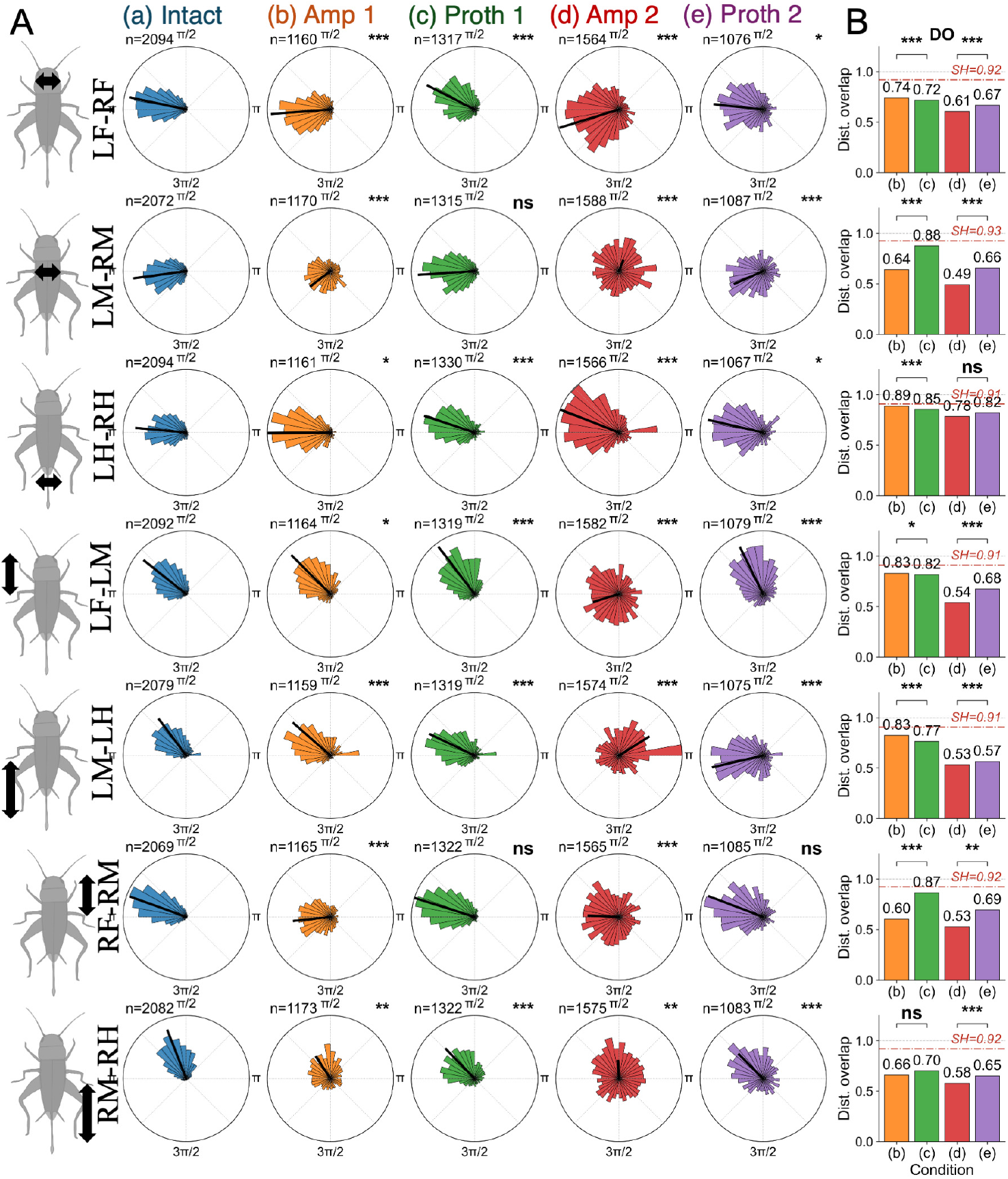
Circular distributions of inter-leg phase differences for all leg-pair combinations across experimental conditions. Rose histograms illustrate the circular distributions of inter-leg phase differences for all seven leg pairs (rows) across five experimental conditions (Intact, Amp 1, Proth 1, Amp 2, Proth 2; columns). Leg pairs are classified by anatomical grouping: bilateral forelegs (LF–RF), bilateral middle legs (LM–RM), bilateral hindlegs (LH–RH), ipsilateral fore–middle pairs (LF–LM, left; RF–RM, right), and ipsilateral middle–hind pairs (LM–LH, left; RM–RH, right). Each histogram comprises 36 equal-width bins (10^◦^ per bin); the radial scale is normalized to the maximum density across conditions to facilitate direct shape comparison, and a radial line indicates the mean resultant vector. Sample size (*n*) is shown in the upper left of each panel, where *n* denotes the number of LH-touchdown events analysed. Statistical comparisons against the Intact condition were performed using the Watson–Williams two-sample test with Benjamini–Hochberg false-discovery-rate (BH-FDR) correction; significance levels are indicated as *p <* 0.05 (*), *p <* 0.01 (**), *p <* 0.001 (***), or non-significant (ns). The rightmost column shows the distribution overlap (DO) between each condition and the Intact reference (DO = 1.0: complete overlap; DO = 0.0: no overlap), quantifying similarity in circular probability-density profiles. In the Proth 1 condition, the right middle leg (RM) carries the prosthetic limb; pairs involving RM are intact–prosthetic (LM–RM, RF–RM, RM–RH), whereas all remaining pairs are intact–intact. Notably, intact–intact pairs not directly involving the prosthetic limb (RF–L_1_F_2_and LF–LM) also exhibit statistically significant phase-difference changes in Proth 1, indicating that prosthetic integration induces network-wide coordination reorganisation extending beyond the directly substituted segment (see Discussion). In each DO panel, the red dash-dot line denotes the split-half distribution overlap (SH-DO) for the Intact condition, computed between odd- and even-indexed stride cycles, and provides an empirical reference for within-condition distributional variability.

For forward and angular velocities in each condition (Fig. 3), we assessed statistical differences relative to the Intact condition using two-sided Mann–Whitney U tests. Raw *p*-values were corrected for multiple comparisons across conditions using the Benjamini–Hochberg false discovery rate (FDR) procedure. Only the significance category (ns, *, **, ***) was displayed in Fig. 3, while the numerical *p*-values were reported in Table S2.

**Figure 3:**
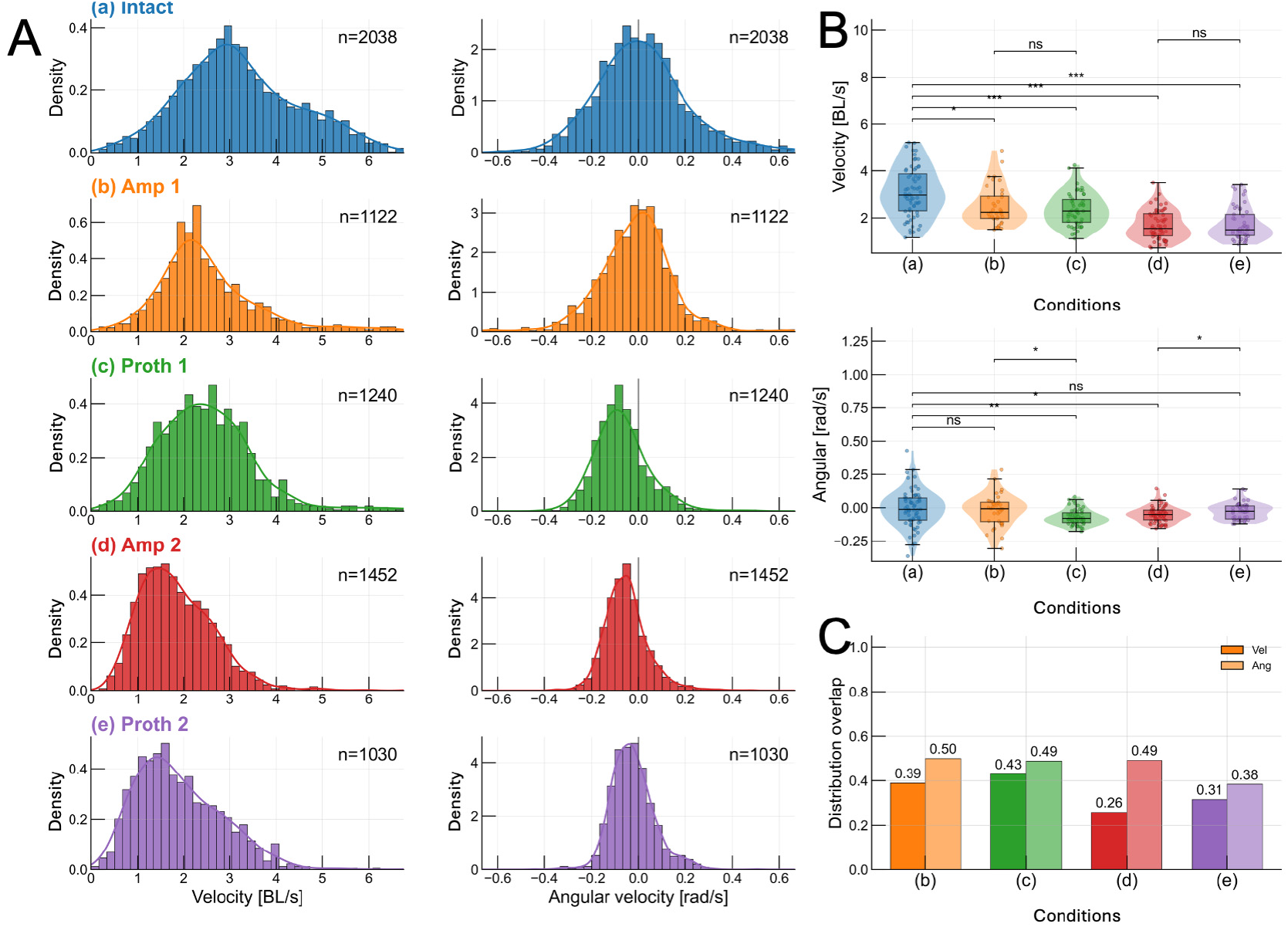
Condition-dependent Changes in Normalized Forward and Angular Velocities. (**A**) Normalized forward velocity (BL/s) and angular velocity (rad/s) distributions across five conditions: (a) Intact, (b) Amp 1, (c) Proth 1, (d) Amp 2, and (e) Proth 2. Histograms (filled bars) are overlaid with Gaussian kernel density estimates, with sample sizes (*n*) indicated in each panel. (**B**) Top right: velocity comparison: violin plots with overlaid box plots and jittered points summarize the distribution of median forward velocity across the five conditions (a)–(e). Middle right: angular velocity comparison: same representation as the top right panel. In both summary panels, statistical differences are assessed between the Intact condition and each manipulation (Amp 1, Proth 1, Amp 2, Proth 2) utilizing a two-sided Mann–Whitney U test, followed by Benjamini–Hochberg false discovery rate correction across all pairwise comparisons. Corrected *p*-values are reported in Table S2, with only the corresponding significance levels indicated on the plots (ns, *, **, ***). (**C**) Bottom right: distribution overlap (DO): the similarity between the Intact distribution and each manipulated condition is quantified as a histogram-based distribution overlap for forward velocity (darker bars) and angular velocity (lighter, same-hue bars). Higher DO values indicate greater preservation of the intact kinematic distribution under each manipulation.

Spatial foot-placement distributions were compared between each manipulation condition and the Intact condition using the two-sample Kolmogorov–Smirnov test, applied separately to the lateral (*x*) and anteroposterior (*y*) coordinates of each leg; both the significance and the effect size (the *D* statistic) were recorded. To preserve independence, foot placements were sampled once per stride cycle at the anteroposterior excursion peaks of each leg rather than frame-by-frame. The resulting *p*-values were corrected for multiple comparisons across all leg–axis–condition combinations using the Benjamini–Hochberg false-discovery-rate procedure. Because the stride-level sample sizes are large (*n* ≈ 1,300–2,600), the test is expected to reach significance even for small distributional differences. Recovery was therefore assessed based on the magnitude of deviation, quantified by the *D* statistic and the distribution overlap relative to the intact split-half (SH-DO) reference, rather than on significance alone.

### 2.7 Distribution Overlap (DO) Analysis

To quantify the similarity between the intact condition and the manipulated conditions (b)–(e), we employed the distribution overlap (DO), a bounded measure of probabilistic similarity [48, 49]. For two normalized histograms with bin masses {*p_k_*} and {*q_k_*} (Σ*_k_ p_k_* = Σ*_k_ q_k_* = 1),

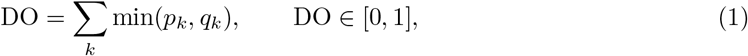

where DO = 1 indicates identical distributions and DO = 0 completely non-overlapping ones.

For the inter-leg phase differences, which are circular quantities, DO was evaluated on the periodic domain *ϕ* ∈ [0, 2*π*) rather than on a linear one. The domain was partitioned into *K* = 36 equally spaced bins of width 2*π/K* (i.e. 10^◦^), *B_k_* = [2*π*(*k* − 1)*/K,* 2*πk/K*) for *k* = 1*, . . . , K*. Because the partition spans the full period with *B_K_* adjacent to *B*_1_, the periodic boundary condition *ϕ* = 0 ≡ *ϕ* = 2*π* is satisfied without any boundary discontinuity. For the forward and angular velocities and the foot trajectories, the same estimator was applied to linear histograms computed over a common data range, using 40 bins for the velocities and 100 bins for each of the fore–aft (*x*) and vertical (*y*) trajectory components. For each condition, DO was computed separately relative to the intact distribution. In Fig. 3C, forward-velocity DO is drawn in the base condition color and angular-velocity DO in a perceptually lightened hue of the same color.

To provide an empirical, data-derived reference for the absolute interpretation of DO, we additionally computed a split-half DO (SH-DO) for the intact condition. Intact observations were divided into two non-overlapping subsets by alternating odd- and even-indexed stride cycles for the phase differences (Fig. 2) and alternating odd- and even-indexed trials for the foot trajectories (Fig. 4). DO was then computed between the two subsets. The SH-DO quantifies the distributional reproducibility expected between independent subsamples of unperturbed locomotion, and thereby provides a baseline for judging whether a condition falls within the range of natural within-condition variability. It is shown as a red dash-dot line in the DO panels of Figs. 2 and 4.

**Figure 4:**
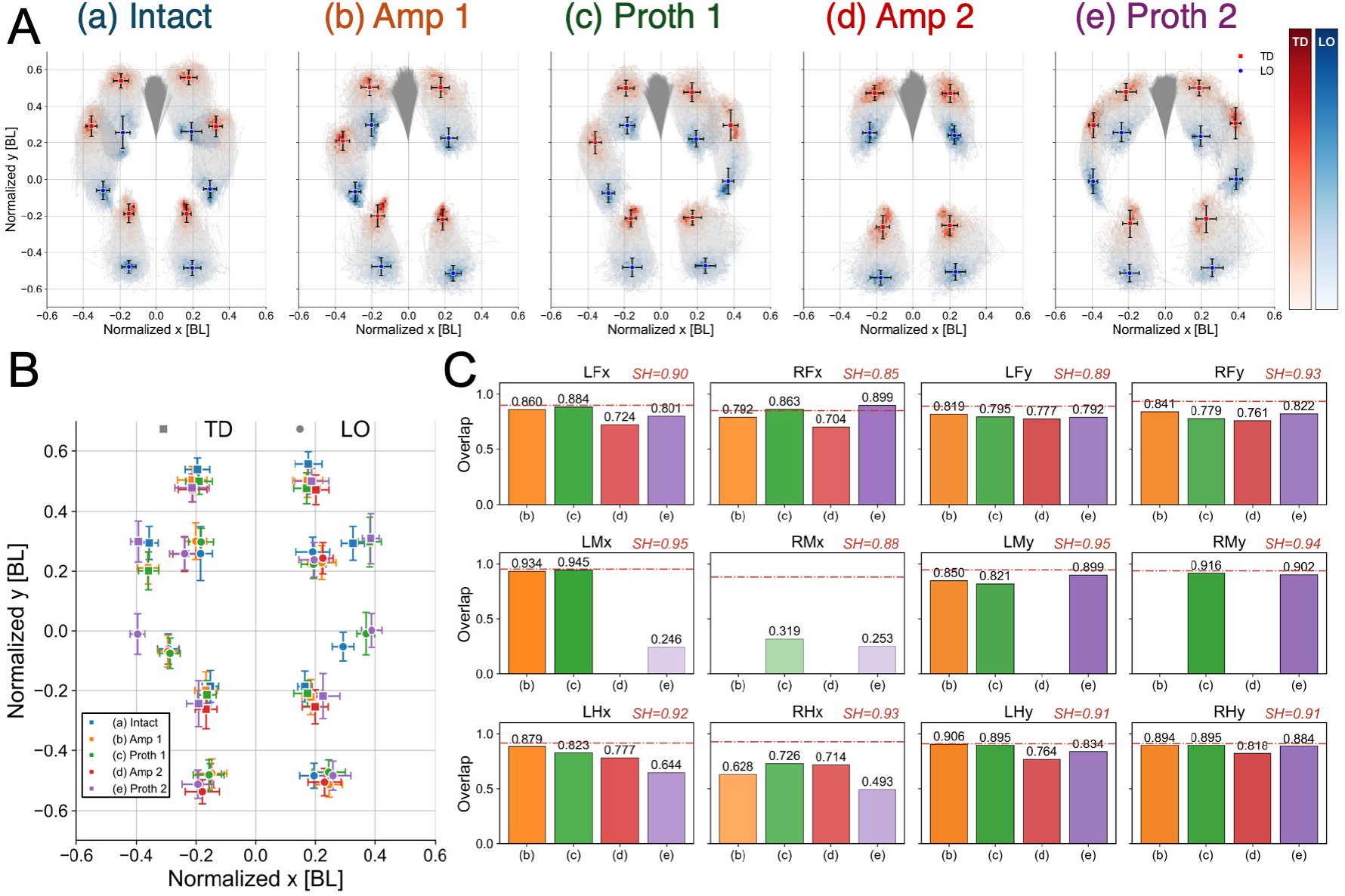
Spatial Distributions and Mean Touchdown (TD) and Lift-off (LO) Positions of the Fore–mid–hind Leg Tarsi across Five Conditions. (**A**) Density maps represent the normalized 2D spatial probability of TD (red) and LO (blue) events aggregated across individuals. Warmer red and blue colors indicate higher event density. Superimposed trajectories of selected body landmarks (head-pronotum segments, LF, LM, LH, RF, RM, and RH leg tarsi) illustrate the spatial envelope of body motion in the body coordinate system. Mean TD positions (red squares) and mean LO positions (blue circles) are plotted with standard-deviation (SD) error bars for each leg. Columns correspond to the conditions (a)–(e) in the Fig. 1, respectively. Axes represent body-length–normalized coordinates (BL), allowing direct comparison across individuals. (**B**) Comparison of the averaged TD and LO positions with SD expressed in body-length–normalized coordinate across all experimental conditions. Squares and circles indicate TD and LO events, respectively, while colors encode experimental conditions. (**C**) Distribution overlap of foot trajectories along the fore–aft (*x*) and vertical (*y*) axes across experimental conditions. Quantitative distribution overlap (DO) between the intact condition and each experimental condition (b)–(e) is shown for all six legs. For each leg, DO values were computed independently for the *x*- and *y*-axes using normalized one-dimensional trajectory distributions. Bar transparency scales with DO (higher DO indicates greater similarity to intact). In each DO panel, the red dash-dot line denotes the split-half distribution overlap (SH-DO) of the Intact condition, computed between odd- and even-indexed trial recordings, and provides an empirical reference for within-condition variability in foot-trajectory position.

For the phase differences, DO and the Watson–Williams test capture complementary, non-redundant aspects of coordination: DO quantifies the similarity of distributional *shape* (spread, modality, and overall overlap), whereas the Watson–Williams test evaluates whether the *mean phase direction* differs. Because these are independent dimensions, high DO can co-occur with a significant Watson–Williams result (similar shape but a shifted mean) and low DO with a non-significant one (wider spread but a preserved mean). We therefore operationally defined recovery as simultaneously satisfying both criteria: a DO value at or above the SH-DO reference and no significant difference in mean phase direction (*p >* 0.05). Conditions meeting only one criterion were classified as showing partial recovery.

## 3 Results

### 3.1 Reorganization of Inter-Leg Phase Coupling Under Amputation and Prosthetic Manipulations

To quantify changes in inter-leg coordination across leg amputation and prosthetic manipulations, we analyzed the circular distributions of inter-leg phase differences at the touchdown timing of the LH leg under five experimental conditions (Intact, Amp 1, Proth 1, Amp 2, and Proth 2). Fig. 2 presents the complete phase-difference analysis for all leg-pair combinations, classified by the functional status of each limb (intact–intact, intact–prosthetic, or prosthetic–prosthetic). In the Intact condition, all leg pairs exhibited sharply concentrated unimodal phase distributions, as indicated by long mean resultant vectors, reflecting the stable and stereotyped phase relationships characteristic of tetrapod gait in insect locomotion.

Leg amputation produced condition-specific distortions in these distributions. Both the Amp 1 and Amp 2 conditions resulted in moderate distributional broadening and, for certain leg pairs, subtle shifts in the preferred phase; for example, the RF–RM phase difference shifted toward *π*. These changes were accompanied by shorter resultant vector lengths and increased phase-angle variability. By contrast, the prosthetic conditions (Proth 1 and Proth 2) exhibited reduced phase-difference variability, indicating recovery toward distributions that more closely resembled those observed in the Intact condition. The Watson–Williams test (with BH-FDR correction) (detailed in Table S1) identified significant alterations in the circular mean for the leg amputation conditions relative to the Intact condition (*p <* 0.05 or *p <* 0.01). However, no significant differences were detected for the Proth 1 condition in the RM–LM phase difference or for the Proth 1 and Proth 2 conditions in the RF–RM phase difference, indicating statistically equivalent recovery of these phase relationships relative to the Intact condition. Direct Watson–Williams tests (BH-FDR corrected across all 42 pairwise tests) between corresponding amputation and prosthetic conditions revealed significant differences in inter-leg phase coordination for six of seven leg pairs in both comparisons: Amp 1 vs. Proth 1 showed significant differences in LF–RF, LM–RM, LH–RH, LF–LM, LM–LH, and RF–RM (all *p*_adj_ *<* 0.001, ***; LF–LM: *p*_adj_ = 0.015, *), with RM–RH non-significant (*p*_adj_ = 0.065, ns); Amp 2 vs. Proth 2 showed significant differences in LF–RF, LM–RM, LF–LM, LM–LH, and RM–RH (all *p*_adj_ *<* 0.001, ***; RF–RM: *p*_adj_ = 0.0047, **), with LH–RH non-significant (*p*_adj_ = 0.064, ns). These results provide direct statistical evidence that prosthetic integration altered inter-leg phase relationships relative to the amputated state in six of seven leg pairs in each prosthesis application. Notably, in the Proth 1 condition, intact–intact leg pairs not directly involving the prosthetic limb, including the bilateral foreleg pair RF–LF and the ipsilateral fore–middle pair LF–LM, also exhibited significant changes in phase difference relative to the Intact condition (Watson–Williams test with BH-FDR correction, *p <* 0.05). This finding indicates that prosthetic integration induced coordination changes beyond the directly substituted segment, consistent with inter-segmental signal propagation within the distributed CPG network [12, 50]. Furthermore, in the Proth 2 condition, an asymmetry was observed between the left and right sides: LF–LM showed a significant phase shift whereas RF–RM did not, a pattern we interpret as a history-dependent consequence of the sequential within-subject experimental design, in which the right middle leg underwent one additional adaptation cycle before bilateral prosthetic integration.

To further quantify distributional similarity, we computed the distribution overlap (DO) between each condition and the Intact baseline; we additionally computed the split-half DO (SH-DO) of the Intact condition by partitioning odd- and even-indexed stride cycles and computing DO between the two subsets, providing a data-derived reference for within-condition variability (red dash-dot line in Fig. 2; see Materials and Methods). DO values systematically decreased with leg amputation (Amp 1 and Amp 2) but exhibited recovery with the introduction of prostheses (Proth 1 and Proth 2), suggesting that phase distributions approached those of the Intact condition. For several leg pairs, the DO values under Proth 1 and/or Proth 2 reached or exceeded the SH-DO reference, indicating recovery to within the range of natural within-condition variability. This finding implies a significant reconstruction of the underlying phase-coupling dynamics with the use of prostheses. Together, these patterns indicate that prosthetic limb integration does not simply restore the original inter-leg coordination topology but rather induces network-level reorganisation that reaches a new functional equilibrium, as discussed below. Notably, this temporal recovery occurred despite the prosthetic limb’s substantially greater mass and complete lack of articulated distal joints. As established in the Discussion, these mechanical properties do not compromise campaniform sensilla activation because CS responds to load onset at ground contact rather than to joint kinematics [51].

### 3.2 Reorganization of Locomotor Velocity Distributions Under Amputation and Prosthetic Manipulations

Leg amputation and prosthetic leg implementation resulted in state-dependent distortions in the characteristics of forward and angular velocity (Fig. 3). In intact subjects, forward velocity displayed a bimodal profile, characterized by a velocity-dependent transition in gait (details provided in Fig. S5), while angular velocity exhibited a broader distribution compared to other conditions. The Amp 1 condition caused a significant shift in the velocity distribution toward lower values, whereas the angular velocity distribution showed asymmetry due to morphological discrepancies from amputation at the FTi joint of the RM leg. In the Proth 1 condition, where a prosthetic leg was added to the FTi joint of the RM leg, there was a modest recovery in the velocity distribution toward faster ranges. The Proth 1 condition resulted in a significantly shifted angular velocity distribution relative to the Intact condition (*p*_adj_ = 0.0035, Mann–Whitney *U* , BH-FDR corrected), reflecting a consistent directional bias attributable to the structural asymmetry introduced by unilateral prosthetic attachment (see Fig. S10). Under the Amp 2 condition, in which both the LM and RM legs were amputated at the FTi joint, the velocity distribution shifted further toward lower velocities, while the angular velocity distribution recovered a more symmetrical pattern, although it remained imperfectly symmetrical, due to bilateral amputation. Finally, in the Proth 2 condition, which involved attaching prosthetic legs to both the LM and RM legs, we observed a recovery of the velocity distribution toward the intact state compared to the Amp 2 condition, with the angular velocity distribution also approaching a symmetrical pattern. To assess heading direction across conditions, we computed the cumulative angular displacement per trial (stride-by-stride integral of angular velocity); unilateral amputation (Amp 1) produced no significant directional drift relative to the Intact condition, whereas the Proth 1 and Amp 2 conditions showed a consistent directional bias that was partially resolved upon bilateral prosthetic integration in the Proth 2 condition (Fig. S10).

Quantitative analysis of median velocities supported these trends (Table S2). Median forward velocity was significantly reduced across all conditions compared to the intact state, as indicated by Benjamini–Hochberg false discovery rate (FDR)-corrected *p* values (significance is depicted in the figure). Regarding angular velocity, the Proth 1 condition resulted in a significantly asymmetric angular velocity distribution due to structural asymmetry between the left and right sides. Conversely, under the Proth 2 condition, the addition of prostheses to both LM and RM legs did not result in a significant difference relative to the intact distribution, suggesting a recovery to a symmetric angular velocity distribution. Direct pairwise comparisons (Mann–Whitney *U* test, BH-FDR corrected across all 12 tests) showed that locomotion velocity did not differ significantly between the amputation and corresponding prosthetic conditions (Amp 1 vs. Proth 1: *p*_adj_ = 0.713, ns; Amp 2 vs. Proth 2: *p*_adj_ = 0.815, ns), while angular velocity was significantly higher in the prosthetic conditions (Amp 1 vs. Proth 1: *p*_adj_ = 0.021, *; Amp 2 vs. Proth 2: *p*_adj_ = 0.041, *; see Fig. 3).

To assess the extent to which each manipulation preserved the overall structure of intact forward and angular velocity characteristics, we computed a distribution-overlap (DO) metric between the intact distribution and each manipulated condition. Consistent with prior results (Fig. 2 B), the forward velocity distribution exhibited distinct changes due to the amputation manipulation and a corresponding recovery to the intact distribution following the prosthesis manipulation. For the angular velocity distribution, the Proth 2 condition displayed the most pronounced differences in distribution similarity relative to the intact condition (notably, DO was not included in the assessment of distribution asymmetry).

The partial recovery of locomotion speed following prosthetic attachment is consistent with the CPG re-entrainment demonstrated in Section 3.1: restoration of campaniform sensilla-mediated load-onset signals re-engages inter-ganglionic phase coupling, and the consequent normalisation of CPG step frequency is expected to promote speed recovery [50]. However, complete speed recovery was constrained by the prosthetic limb mass (approximately 8× that of the intact middle leg), which imposes an elevated inertial load during swing phase and limits the achievable step frequency. This dissociation, characterized by full recovery of temporal coordination despite incomplete recovery of speed, provides additional evidence that inter-leg phase regulation and overall locomotion speed are governed by overlapping but mechanistically distinct processes. The former requires only the restoration of load-onset signals, whereas the latter additionally requires mechanical optimization of limb inertia.

### 3.3 Reorganization of Spatial Distributions of Fore–Mid–Hind Leg Tarsi Positions Under Amputation and Prosthetic Manipulations

Across all animal subjects, the tarsal trajectories of the fore-, middle-, and hind-legs, along with touchdown (TD) and liftoff (LO) events, exhibited highly structured spatial distributions closely aligned with whole-body kinematics (Fig. 4 A). As illustrated in Fig. 4 A (a), the Intact condition showed a structured and highly symmetric spatial coordination of foot trajectories, serving as a reference pattern for normal locomotion. The averaged spatial distributions for TD and LO within each condition (Fig. 4 B) revealed six well-defined, leg-specific regions in body-length–normalized coordinates. These regions highlighted systematic spatial deviations from the typical coordinated pattern of the fore-, middle-, and hind-legs observed during stable intact walking.

In the (b) Amp 1 condition (orange), the amputation of the RM leg resulted in significant alterations in the TD and LO locations of the ipsilateral fore- and hind-legs. Notably, the RF leg exhibited a posterior shift in TD, along with both posterior and laterally outward shifts in LO. The RH leg demonstrated a posterior and laterally outward shift in LO, while its TD position remained largely unchanged. In contrast, the LF leg experienced a reduction in step length, characterized by a posterior shift in TD and an anterior shift in LO, whereas the LM leg exhibited only a posterior shift in TD. No substantial changes were detected in the TD or LO positions of the LH leg. In the (c) Proth 1 condition (green), an artificial prosthetic leg was affixed to the FTi joint of the amputated RM leg, facilitating the participation of the prosthesis in gait pattern generation. Due to the absence of articulated joints in the prosthesis, its foot trajectory did not fully replicate that of the Intact condition. Nonetheless, the anteroposterior (*y*-axis) range of the trajectory, defined by the mean TD and LO positions, was comparable to that of intact walking. Despite marked phase recovery (Fig. 2), the spatial recovery of the TD-LO event distribution was limited, except for a partial restoration in the lateral (*x*-axis) positions of the fore-legs. In the (d) Amp 2 condition (red), bilateral amputation of the middle legs resulted in symmetrical trajectory changes in both the fore-and hind-legs, mirroring the unilateral effects observed in (b) Amp 1 but affecting both sides. The forelegs exhibited posterior shifts in TD and posterior and laterally outward shifts in LO, while the hind legs displayed similar posterior and laterally outward shifts in LO, with the TD positions of the hindlegs also displaced posteriorly. In the (e) Proth 2 condition (purple), bilateral prostheses were attached to the middle legs. Consistent with (c) Proth 1, the anteroposterior TD–LO ranges of the prosthetic middle legs closely matched those observed in the Intact condition. In terms of spatial coordination, partial recovery was noted in the lateral (*x*-axis) positions of the forelegs and the anterior (*y*-axis) positions of the hind legs. In contrast, the remaining spatial distributions largely maintained the modified patterns characteristic of the (d) Amp 2 condition.

To quantify the impact of each manipulation on the spatial distribution of leg trajectories relative to intact locomotion, we calculated the distribution overlap (DO) of foot trajectories along the lateral (*x*) and anteroposterior (*y*) axes for all six legs (Fig. 4 C). This metric evaluates the extent to which a trajectory retains the statistical spatial structure of intact locomotion, with higher DO values indicating greater similarity. The SH-DO reference (red dash-dot line in Fig. 4), computed between odd- and even-indexed Intact trials, confirmed that the foot-trajectory distributions in the prosthetic conditions remained substantially below the level of natural within-condition variability, providing a data-derived criterion for the persistent spatial deficit.

Consistent with the previously presented spatial analyses, the DO values of the bilateral prosthetic middle legs exhibited a significant directional asymmetry. Although the prostheses did not replicate the lateral (*x*-axis) trajectory distributions observed under the Intact condition, their anteroposterior (*y*-axis) distributions closely mirrored those of intact locomotion, supporting the qualitative spatial patterns discussed earlier. Two-sample Kolmogorov–Smirnov tests (BH-FDR corrected; Supplementary Table S4) showed that every manipulation condition differed significantly from the Intact condition for all legs and axes (*q <* 0.001), reflecting the high sensitivity of the test at the large stride-level sample sizes analysed here. The effect sizes localise the deficit: the lateral foot placement of the prosthetic middle legs showed the largest deviations (e.g. Proth 1 RM, *D* = 0.60, DO = 0.32 vs. SH-DO = 0.88), whereas their anteroposterior placement remained close to the intact reference (*D* = 0.19, DO = 0.92), quantitatively corroborating this incomplete and spatially selective recovery of foot-placement kinematics. For the foreleg pair (LF and RF legs), DO values for both lateral (*x*) and anteroposterior (*y*) distributions remained consistently high (*>* 0.70) across all experimental conditions. This consistency suggests a significant preservation of foreleg trajectories despite amputation and prosthetic interventions. Notably, the transition from Amp to Proth conditions quantitatively indicated a recovery of the lateral (*x*-axis) spatial distribution structure of foreleg trajectories. In contrast, for the hindlegs (LH and RH legs), the similarity in lateral (*x*-axis) trajectory distributions diminished in both Amp and Proth conditions compared to the Intact condition. However, along the anteroposterior (*y*) axis, the hindlegs showed a marked recovery in DO values when transitioning from Amp to Proth conditions, signifying a partial restoration of fore–aft spatial coordination despite ongoing lateral deviations.

The incomplete recovery of foot-trajectory kinematics provides the spatial counterpart to the temporal recovery demonstrated in Section 3.1, jointly establishing the spatiotemporal dissociation that constitutes the central finding of this study. Spatial coordination in insect locomotion depends critically on continuous joint-angle feedback from the femoral chordotonal organ (fCO), which encodes angular position and velocity throughout swing phase to regulate foot placement [14, 52]. Because the prosthetic limb lacks articulated joints, fCO activity cannot be re-established upon prosthetic attachment. The persistent spatial deficit therefore follows *a priori* from the differential anatomy of the campaniform sensilla, which are proximal, contact-activated, and restored by the prosthesis, and the femoral chordotonal organ, which is distal, movement-activated, and not restored by the prosthesis. This mechanistic prediction is confirmed by the present kinematic data and is developed further in the Discussion.

## 4 Discussion

Here, we systematically examine the effects of leg amputation and body-integrated prosthetic legs on the coordinated walking of insects, especially to elucidate the efficacy of prosthetic legs in restoring motor coordination across various dimensions of walking dynamics. The present study reveals a hierarchical, sensor-specific recovery of coordinated locomotion following prosthetic limb integration in crickets. At the first tier, inter-leg phase coordination, governed by the CPG network and entrained by campaniform sensilla (CS) load signals, fully recovers upon prosthetic attachment (Section 3.1). The prosthetic limb contacts the substrate, generates ground reaction forces, and thereby re-establishes the CS input required for CPG re-entrainment. At the second tier, walking speed partially recovers as the functional consequence of restored CPG output, but is attenuated by the elevated inertial load of the prosthesis (Section 3.2). At the third tier, spatial foot-placement precision does not recover (Section 3.3): the wire-frame prosthesis lacks articulated joints, precluding activation of the femoral chordotonal organ (fCO) that encodes swing-phase limb position, and thus leaving proprioceptive spatial guidance absent. This hierarchical structure directly reflects the sensor-specific dependence on joint kinematics established below: CS activation requires only ground contact (restored by the prosthesis), whereas fCO activation requires joint movement (not restored). The resulting pattern of selective recovery of temporal coordination without recovery of spatial coordination constitutes the spatiotemporal dissociation that is the central finding of this study.

Leg amputation in insects disrupts not merely a propulsive appendage but the sensory-motor loops that support inter-leg coordination, particularly those conveying information on leg loading, unloading, and joint movement [50, 53]. In intact insects, the legs are densely equipped with campaniform sensilla encoding force and load transfer [51, 53–56], together with chordotonal organs and other proprioceptors encoding joint kinematics [50, 57], providing continuous feedback to thoracic circuits that dynamically adjust inter-leg phase relationships and motor output [52, 58]. Amputation removes these sensory streams and disrupts load-dependent phase coupling between adjacent legs, a process relying critically on mechanically grounded feedback rather than centralized timing alone [59, 60], thereby reducing walking speed and altering turning behavior and muscle activation [25, 61]. The partial restoration of coordinated walking following prosthetic integration should not be interpreted as a replacement for central rhythm generation, but as the re-establishment of mechanically relevant sensory feedback that enables the existing neural control architecture to resume coordinated gait production. By re-establishing ground contact and bearing load during stance, the prosthesis likely reinstates the unloading and load-transfer signals that trigger phase transitions and inter-leg handover [52]. Because campaniform sensilla robustly encode changes in force and bending moments [51, 55], these signals remain functionally effective even without precise morphological replication of a biological leg, provided that the key mechanical events of load onset and release are preserved [56, 62]. This is consistent with evidence that peripheral sensory feedback can override intrinsic bilateral synchronizing tendencies in central circuits to stabilize the canonical alternating gait [25], and with observations across insect taxa that locomotor systems self-organize under sensory-mechanical constraints following amputation [11, 50, 63]. Crucially, the comprehensive inter-leg phase analysis (Fig. 2) shows that these effects are not confined to the directly substituted segment but propagate through the distributed CPG network: in the Proth 1 condition, intact–intact leg pairs (RF–LF, LF–LM) exhibited significant phase-difference changes alongside the recovery of the directly coupled intact–prosthetic pairs (LM–RM, RF–RM). This pattern indicates that the reinstated campaniform sensilla load-timing signal from the prosthetic limb re-entrains the middle-leg CPG oscillator, which in turn modifies its coupling with both ipsilateral and contralateral oscillators.

The resulting network state therefore does not represent a simple restoration of the pre-amputation coordination pattern, but rather a new stable configuration that nonetheless supports functional locomotion. This reflects motor degeneracy, in which multiple inter-leg phase topographies can sustain adequate locomotor performance [64, 65]. The asymmetric partial recovery observed in Proth 2 (LF–LM significant, RF–RM not) further reflects the history dependence of this adaptation, as the sequential within-subject design causes the left and right middle-leg oscillators to approach their new stable states along different trajectories, thereby producing left–right asymmetry despite bilaterally symmetric prosthetic conditions. These patterns extend the spatiotemporal dissociation framework and remain mechanistically consistent with the CS–fCO asymmetry established above: prosthetic reintegration restores temporal coordination in the directly coupled pairs while simultaneously inducing broader coordination reorganisation across the network. Our findings therefore suggest that prosthesis-induced recovery of walking arises from the reinstatement of sensory constraints within the inter-leg coordination network, enabling the locomotor system to reconstruct functional gait patterns through embodied sensory feedback and network-level reorganisation rather than through direct modification of central motor rhythms.

The asymmetric spatial recovery observed in this study, in which the middle-leg prosthesis preferentially restored the lateral (*x*) component of foreleg trajectories while mainly supporting anterior–posterior (*y*) recovery in hindleg trajectories, closely aligns with established biological principles of inter-leg coordination in insects [12, 66]. Walking in insects is organized through a distributed control architecture, with each leg governed by local sensorimotor mechanisms coupled to neighboring legs via state-dependent coordination signals [67–70]. In this framework, the middle legs serve as reference limbs for body support, constraining the kinematics of both anterior and posterior legs [58, 71]. This functional differentiation, with forelegs specialized for exploration and steering, hindlegs for propulsion, and middle legs for load-bearing support, provides the biological basis for Walknet-inspired coordination rules, in which global gait patterns emerge from local inter-leg interactions rather than centralized control [67, 69]. In this context, preferential recovery of the foreleg’s lateral component following middle-leg prosthetic integration suggests that re-established load-bearing and ground contact provided a stable mechanical and sensory reference for body stabilization and steering, enabling more accurate lateral targeting of the forelegs [72, 73]. Conversely, pronounced recovery of the hindleg’s anterior–posterior component is consistent with load-based coordination mechanisms, wherein load transfer and unloading signals from the middle leg regulate stance-to-swing transitions and propulsive force generation [51, 52]. Crucially, this recovery pattern cannot be explained by reflex pathways alone; it aligns with a contemporary view in which segmentally organized CPGs within the thoracic ganglia are continuously modulated by proprioceptive and load-related sensory feedback while being mutually coupled across segments [74, 75]. Experimental evidence shows that leg stepping can entrain neighboring CPGs [76], that thoracic connective lesions profoundly alter inter-leg coordination [77], and that isolated middle-leg preparations retain pattern-generating and sensory-modulated motor capabilities [78, 79]. This supports the interpretation that the middle leg occupies a privileged position in shaping inter-leg coordination dynamics. These findings indicate that the prosthesis did not merely compensate for lost limb geometry but selectively reinstated sensory–mechanical constraints along distinct spatial axes, enabling the locomotor system to reassemble functional gait patterns through coordinated interactions between load-dependent feedback, local pattern-generating networks, and intersegmental coupling.

The selective recovery of temporal inter-leg coordination in the presence of only partial recovery of spatial foot placement can be understood within a mechanistic framework in which temporal and spatial coordination depend on functionally distinct sensory feedback pathways. Temporal coordination, defined by the inter-leg phase relationships that determine gait pattern, is governed primarily by intersegmental CPG coupling and entrained by mechanosensory signals encoding the timing and magnitude of ground contact forces during the stance phase [50, 52]. Campaniform sensilla (CS) located in the trochanteral region of each leg are the primary encoders of these load-relevant signals, detecting force onset, magnitude, and inter-leg load transfer [51]. Critically, CS-mediated load signals reflect substrate contact forces at the proximal leg and are transmitted regardless of distal leg geometry [55]. Because the prosthetic leg contacts the substrate during stance and generates ground reaction forces, CS-mediated load-timing signals are plausibly re-established with each prosthetic step cycle, reinstating the timing cues that entrain CPG coupling across thoracic segments. Spatial coordination, defined as the accurate placement of individual feet relative to body position during the swing phase, requires continuous proprioceptive feedback on joint angles throughout swing, encoded primarily by the femoral chordotonal organs (fCO) [14, 80]. These mechanoreceptors encode angular position and velocity of the tibia–femur joint continuously during swing, providing the mid-course corrections necessary for precise spatial targeting of the foot. The passive prosthetic leg used in this study lacks articulated joints; fCO-analogous joint-angle feedback is therefore not reinstated, leaving the continuous kinematic feedback required for spatial foot placement accuracy absent. This mechanistic asymmetry between the reinstatement of CS-mediated temporal signals, which can be achieved through substrate contact alone, and the non-reinstatement of fCO-mediated spatial signals, which requires articulated joints, *predicts, a priori*, the selective recovery of temporal rather than spatial coordination. This prediction is confirmed by our data: inter-leg phase relationships recover robustly across all prosthetic conditions, whereas foot placement trajectories recover only partially and axis-specifically. This concordance between mechanistic prediction and observed recovery pattern constitutes the central mechanistic contribution of this study, and positions the temporal–spatial dissociation as evidence for the hierarchical organization of distributed locomotion control.

The selective recovery of temporal gait coordination following passive prosthetic integration exemplifies the principle of morphological computation [8], according to which functionally relevant behavioral outcomes arise not solely from explicit neural computation but also from interactions among body mechanics, neural control, and the environment. In this study, the mechanical compliance of the prosthetic leg reinstated load-onset and load-release events during stance. These events were sufficient to entrain distributed central pattern generator (CPG) circuits across the thoracic ganglia without requiring active joint control or precise morphological replication of the biological limb. This result suggests that the generation of load-contact signals is a critical functional design requirement for biohybrid locomotor devices: engineering a prosthetic component to produce reliable stance-phase ground-reaction force transients may exploit the self-organizing properties of distributed locomotor networks more effectively than prioritizing morphological fidelity or passive stiffness profiles alone. The robust recovery of temporal coordination, despite the eightfold excess mass of the prosthesis, further indicates that the load-dependent entrainment mechanism can tolerate substantial mechanical perturbations, supporting its potential relevance as a broadly applicable design principle for biohybrid systems.

These findings also have important implications for prosthetic limb design and sensorimotor rehabilitation in vertebrate systems, although the transfer of mechanisms across taxa requires independent validation. The campaniform sensilla, which mediate prosthesis-driven temporal recovery in this study [50]–[55], encode the magnitude and timing of leg-substrate loading during stance. Plantar mechanoreceptors and load-sensitive afferents are considered to perform functionally analogous roles in regulating step timing and inter-limb coordination during human walking. This functional parallel suggests that prosthetic socket interfaces designed to prioritize the transmission of load-onset and load-release timing cues to residual-limb somatosensory afferents may preferentially facilitate the recovery of temporal gait coordination. Correspondingly, rehabilitation strategies that treat temporal inter-limb coupling as an early primary endpoint, before complete spatial accuracy is restored, may capitalize on the relative robustness of load-mediated coordination mechanisms demonstrated here. Such an approach would provide a hierarchically structured and mechanistically grounded framework for sensorimotor rehabilitation following limb loss or locomotor injury.

Although our behavioral data cannot directly distinguish active sensorimotor loop re-engagement from passive mechanical stabilization, three lines of indirect evidence support the interpretation that load-mediated mechanoreceptive signaling, rather than inertial mass effects alone, underlies the observed recovery of temporal coordination. First, the prosthetic legs used in this study are approximately eight times heavier than the biological middle legs. A passive mechanical stabilization model predicts that such mass addition would alter locomotor dynamics uniformly across coordination dimensions. The selective recovery of temporal inter-leg coordination in the absence of equivalent spatial recovery is *inconsistent with this prediction*, and instead suggests that the observed recovery reflects dimension-specific neural processes rather than passive mechanical effects. Second, the trochanteral region through which the prosthetic leg is inserted is adjacent to campaniform sensilla (CS) that encode force onset, magnitude, and inter-leg load transfer during stance [51, 56]. These mechanoreceptors transmit load-relevant signals even when distal leg geometry does not precisely replicate biological morphology [55], because the critical variable for inter-ganglionic CPG coupling is the onset and magnitude of ground reaction forces at the proximal leg, not the kinematic trajectory of the distal limb [50, 52]. The prosthetic leg contacts the substrate during stance and generates ground reaction forces; CS-mediated load signals are therefore plausibly reinstated with each prosthetic stance cycle. Third, Noah et al. (2004) demonstrated that a fully denervated cockroach stump, incapable of transmitting distal sensory signals, still promotes gait recovery when it contacts the substrate [37]. This finding independently implicates substrate contact forces at the proximal leg as the critical variable, providing parallel support for the mechanistic plausibility of our interpretation.

From an interdisciplinary perspective, the present findings integrate classical insect neuroethology [11, 12, 68, 74] with contemporary theories of embodied motor control [9, 81] and bio-inspired robotics [8, 82–84], and recast locomotor recovery as a problem of self-organized behavior emerging from restored sensory–mechanical constraints. Rather than viewing prosthetic intervention as anatomical replacement, our results establish a general principle: coordinated locomotion can recover when task-relevant load-dependent sensory feedback and limb–body coupling are reinstated, even in the presence of substantial morphological asymmetry. Because the sensory modalities mediating prosthesis-driven recovery in this study are conserved across insects such as cockroaches [51, 54] and stick insects [52, 55], the mechanisms uncovered here are inherently transferable, providing a common explanatory framework for locomotor resilience across biological systems, as well as a foundation for adaptive design in robotics and prosthetics. Several aspects of the present study warrant further investigation, not as shortcomings, but as productive avenues for future research. First, the prosthetic legs used in this study were approximately eight times heavier than the intact biological legs, reflecting current technical limitations in fabrication. Nonetheless, crickets demonstrated the ability to sustain walking and re-establish inter-leg phase relationships even under this substantial load, underscoring the robustness of load-dependent coordination mechanisms. Consistent with this reasoning, a mass-matched inert attachment that was identical in mass to the prosthesis but lacked ground-contact geometry did not restore inter-leg phase coordination (Supplementary Movies SM12 and SM13), directly demonstrating that mass addition per se is insufficient to account for the observed recovery of temporal coordination. The Second limitation concerns the ecological validity of the spherical treadmill preparation. The tethered spherical treadmill enables stable, high-frame-rate kinematic tracking under controlled conditions and is a widely adopted tool for quantitative locomotion analysis in insects [43]; however, the tethered configuration constrains translational body movement and may alter the ground reaction force dynamics that drive mechanoreceptive load signals relative to free overground walking. Whether the coordination recovery patterns documented here generalize to untethered walking on natural terrain remains an open question that we identify as an important future direction. Third, prosthetic integration was limited to the bilateral middle legs for experimental tractability. Whether comparable prosthetic integration of the forelegs or hindlegs, which serve distinct biomechanical and sensory roles in hexapod locomotion, produces similar spatiotemporal recovery patterns remains an open question for future investigation. In addition, burst-stepping events, defined as rapid, repetitive contacts that can occur following limb amputation [37, 45, 46], were not excluded using an explicit inter-event-interval criterion. Because burst steps produce approximately uniformly distributed phase angles, any such contamination would lower the DO measure, biasing the analysis against detecting coordination recovery; our conclusions are therefore conservative with respect to this potential confound. A key limitation of the present study is that our interpretation of the recovery mechanism rests on behavioral kinematic analysis alone, without direct electrophysiological measurements of sensory or motor neural activity. Consequently, we cannot definitively distinguish between re-engagement of sensorimotor loops via campaniform sensilla activation and passive mechanical stabilization effects. Electrophysiological recording from thoracic ganglia during prosthetic walking, including extracellular recording of CS afferents and interneurons in the meso- and metathoracic ganglia, represents the most direct test of the proposed mechanism and is therefore a priority for future work. Taken together, these limitations do not diminish the central advance of this study. Instead, they delineate a broader research program grounded in a unifying design rule: robust locomotor coordination does not depend on anatomical completeness or precise kinematic replication, but emerges from embodied sensory–mechanical interaction. By experimentally demonstrating this principle, our work provides a generalizable and quantitative framework for understanding and engineering adaptive locomotion across living and artificial systems.

## Acknowledgments

## Author Contributions

D.O. and H.A. conceived the study, designed and performed the biological experiments, and acquired and analyzed the data. D.O. developed experimental setup, custom software and conducted statistical analyses. D.O. and H.A. interpreted the results. D.O. wrote the original draft of the manuscript. H.A. supervised the research and provided critical intellectual input. Both authors secured funding, contributed to manuscript revision, and approved the final version.

## Data Availability

The data that support the findings of this study are available for peer review via a private Zenodo repository. Detailed descriptions of the dataset contents, structure, and usage are provided in Supplementary Data 1. The data will be made publicly available upon publication.

## Materials and Methods

### Trajectory Reconstruction with Optical Sensors

By using two 2-D optical flow sensor values (*dx*_1_*, dy*_1_*, dx*_2_*, dy*_2_), the rotational angular velocities of the spherical treadmill in the yaw, roll, and pitch directions were calculated using the following equations [25, 43]:

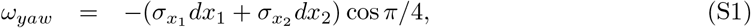

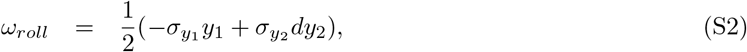

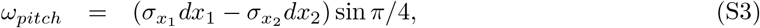

where *σ_x_*_1_ *, σ_y_*_1_ *, σ_x_*_2_ *, σ_y_*_2_ denote the scale parameters for converting millimeter length scale [mm/s] to radian angle scale [rad/s] (see Fig. S1). Next, based on the definition of the coordinate system in the camera frame, the locomotion velocities in the *x* and *y* directions were calculated using the sphere radius (*l_s_* = 50 mm) as follows:

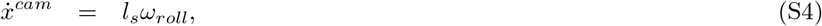

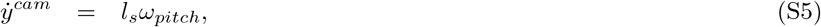

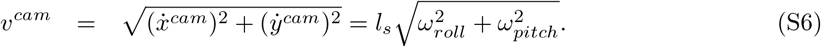

To normalize the length and velocity data based on the body length of crickets, we divided the length and velocity data by the corresponding cricket’s body length (BL) found in Table S3, resulting in a unit transformation from mm or mm/s to BL or BL/s.

The cricket’s walking trajectories were integrated into the camera frame coordinate system by using the velocities from Eqs. (S4) and (S5) with the following equations:

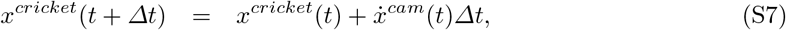

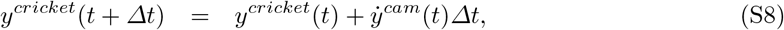

where the initial position is set at the origin, i.e., *x^cricket^*(0)*, y^cricket^*(0) = (0, 0), for each trial.

Caption for Movie SM1. Experimental setup for measuring walking patterns and locomotor velocity in the cricket *Gryllus bimaculatus* with integrated prosthetic legs.

**Caption for Movies SM2 and SM3. Representative walking patterns under the intact condition (a).** Movie SM2 corresponds to a walking speed of 1.71 BL s^−1^ (Fig. S5A), and Movie SM3 to 5.20 BL s^−1^ (Fig. S5B).

**Caption for Movies SM4 and SM5. Representative walking patterns under the Amp 1 condition (b).** Movie SM4 corresponds to a walking speed of 1.08 BL s^−1^ (Fig. S6A), and Movie SM5 to 3.46 BL s^−1^ (Fig. S6B).

**Caption for Movies SM6 and SM7. Representative walking patterns under the Proth 1 condition (c).** Movie SM6 corresponds to a walking speed of 1.83 BL s^−1^ (Fig. S7A), and Movie SM7 to 4.42 BL s^−1^ (Fig. S7B).

**Caption for Movies SM8 and SM9. Representative walking patterns under the Amp 2 condition (d).** Movie SM8 corresponds to a walking speed of 1.01 BL s^−1^ (Fig. S8A), and Movie SM9 to 2.64 BL s^−1^ (Fig. S8B).

**Caption for Movies SM10 and SM11. Representative walking patterns under the Proth 2 condition (e).** Movie SM10 corresponds to a walking speed of 1.96 BL s^−1^ (Fig. S9A), and Movie SM11 to 3.73 BL s^−1^ (Fig. S9B).

**Caption for Movies SM12 and SM13. Representative walking patterns with mass-matched control attachments.** Movie SM12 shows the reversed prosthetic-leg condition, and Movie SM13 shows the straight prosthetic-leg condition. Although both attachments were matched in mass to the functional prosthesis, neither provided appropriate ground-contact geometry, and neither restored inter-leg coordination.

**Caption for Data S1. Synchronized kinematic and locomotion dataset of walking cricket (cricket019).** The dataset comprises two-dimensional *x*–*y* positional trajectories of body and leg markers obtained via DeepLabCut and locomotor velocity signals measured by dual optical flow sensors on a spherical treadmill. The two data streams are temporally aligned using an LED trigger signal. Data and analysis scripts are organized by experimental condition, including intact, leg amputation, and prosthetic leg conditions, and provide the source data for quantitative analyses presented in the main text.

---

**Figure S1:**
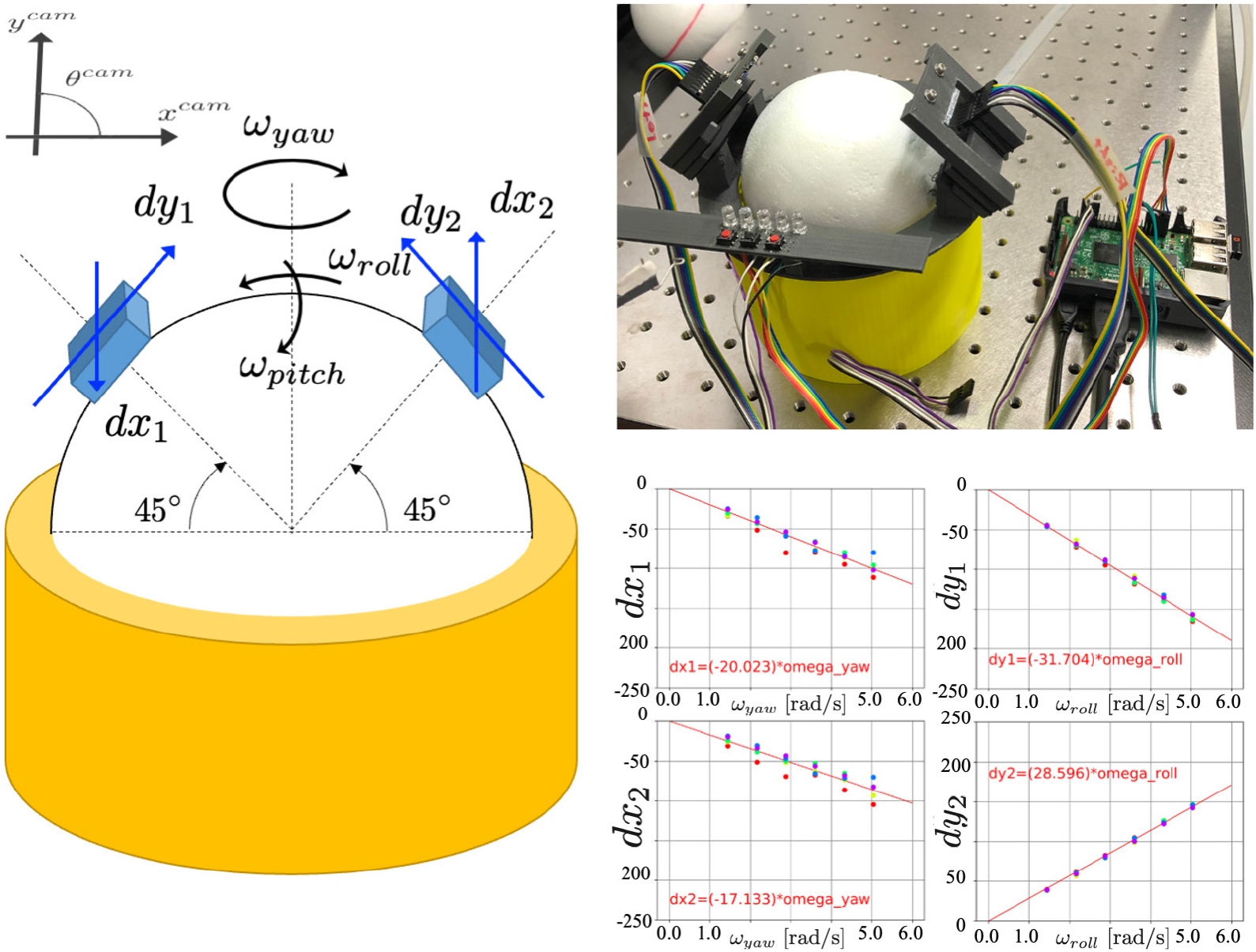
Measurement System for Walking Velocity and Body Rotational Velocity Utilizing Dual Optical Flow Sensors. The left panel illustrates the geometric relationship between the optical flow sensors and the spherical treadmill coordinate system, with the sensors positioned directly above the sphere. The lower-right panel shows the calibration between the angular velocity of spherical rotation and the corresponding sensor outputs. The horizontal axis represents the angular velocities associated with yaw and roll rotations, while the vertical axis denotes the measured sensor values, derived from five repeated measurements at each of six angular velocities. Calibration was conducted using a precisely controlled electric motor (DYNAMIXEL XH430-W210-R, ROBOTIS) to estimate the *σ* parameters in Eqs. S1–S3 [25]. The motor was actuated with continuous, constant-velocity commands via a Raspberry Pi, with the same sphere affixed to the motor to facilitate rotation on the treadmill base. Sensor outputs were recorded during steady spherical rotation in the yaw and roll directions across a range of controlled rotational speeds.

**Figure S2:**
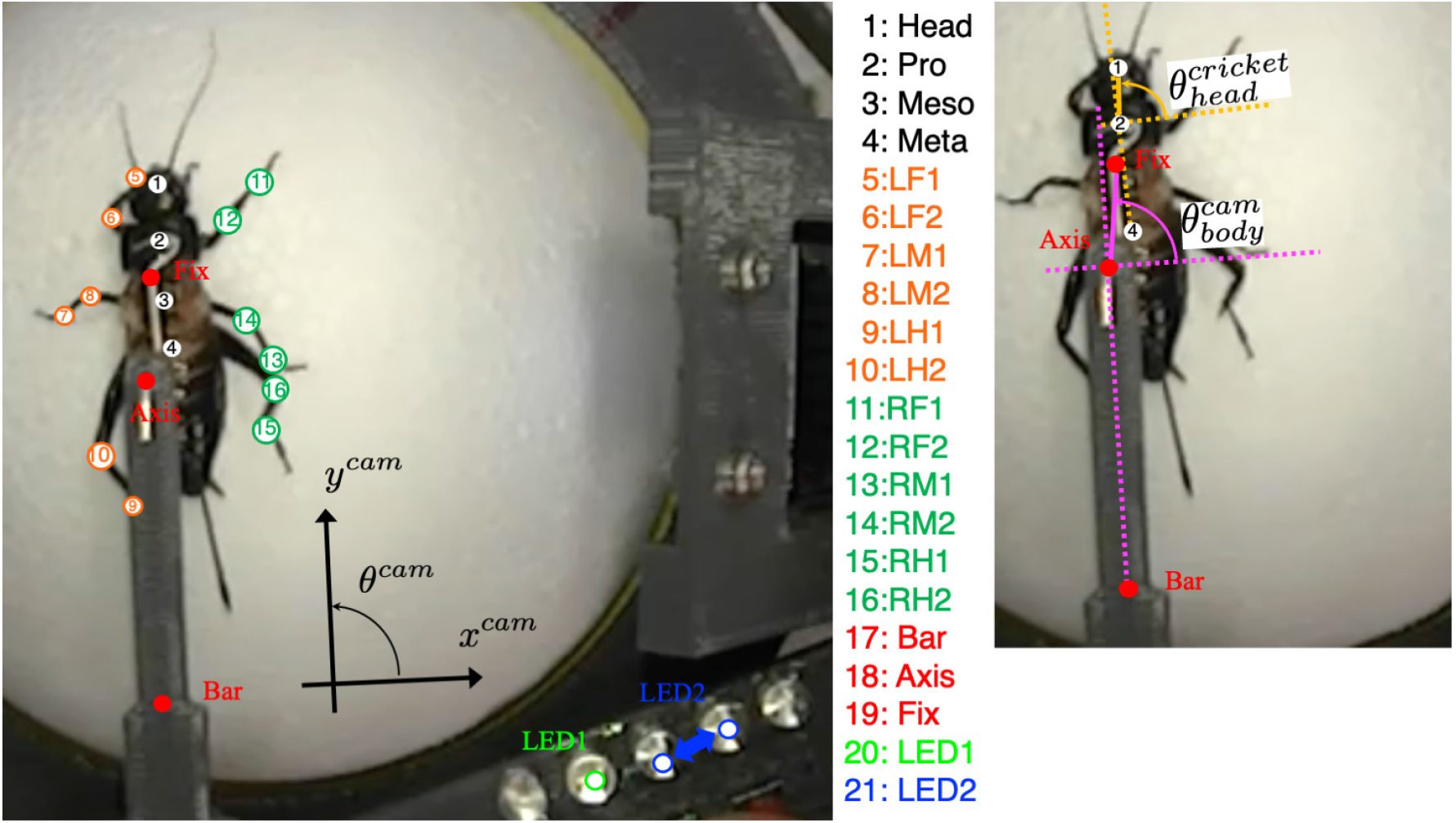
Marker Configuration for Pose Estimation During Walking Using DeepLab-Cut. The positions of markers used for kinematic tracking are indicated, with marker indices displayed at the center of each point. Vertical trajectories of leg joint markers were utilized to delineate power and recovery strokes (stance and swing phases), enabling the quantification of spatiotemporal walking patterns. As shown on the right, head orientation in the cricket reference frame was computed from body landmarks (Head: 1, Pro: 2, Meta: 4), while body orientation in the camera frame was derived from fixed frame markers (Fix: 19, Axis: 18, Bar: 17). The LED2 marker was used to synchronize kinematic data with optical flow sensor recordings.

**Figure S3:**
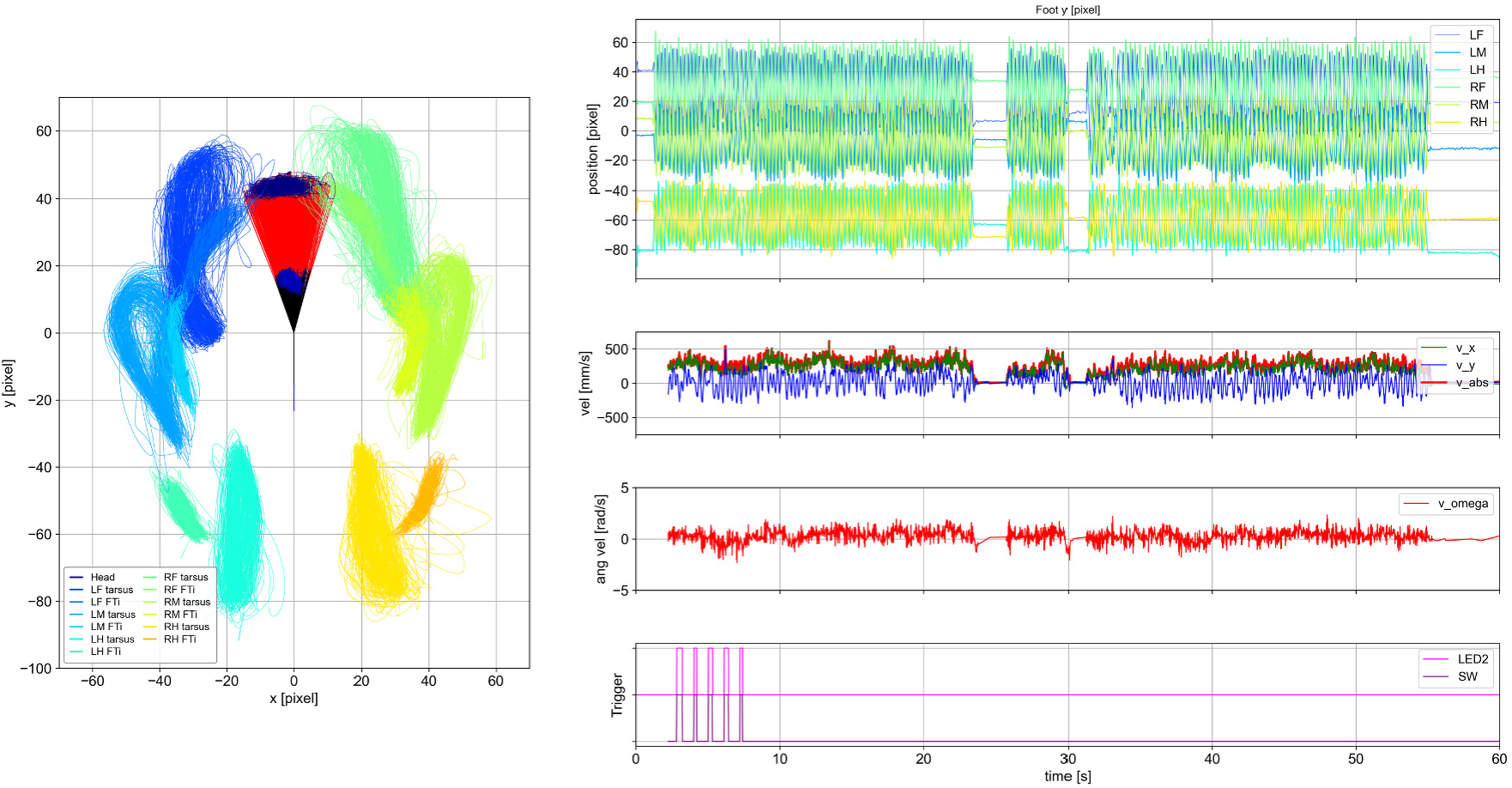
Synchronization of DeepLabCut-Based Kinematics with Walking Velocities Derived from Optical Flow Sensors. The left panel shows *x*–*y* plane trajectories (in pixels) of the Head, Pro, Meso, Meta, LF1, LF2, LM1, LM2, LH1, LH2, RF1, RF2, RM1, RM2, RH1, and RH2 markers extracted using DeepLabCut (DLC). The coordinate system was defined with the Meso marker as the origin, and the y-axis aligned with the line connecting the Meso and Meta markers. The Head–Pro segment is shown in red, and the Pro–Meso segment in black. The right bottom panel illustrates synchronization triggers: the magenta trace represents the DLC-estimated position of LED2 (see Fig. S2), while the purple trace denotes the tactile switch signal for LED2 recorded on the Raspberry Pi. The top panel displays DeepLabCut-estimated vertical leg positions (*y*) for each leg. The plot colors correspond to each leg in the left panel. The right second panel from the top shows forward velocities calculated by the optical flow sensors (green: (*x*), blue: (*y*), red: absolute velocity), and the right third panel illustrates the angular velocity of body rotation about the yaw axis *ω_yaw_* on the treadmill. These plots display the raw data. Trigger-based alignment facilitated the simultaneous analysis of leg kinematics and walking velocity.

**Figure S4:**
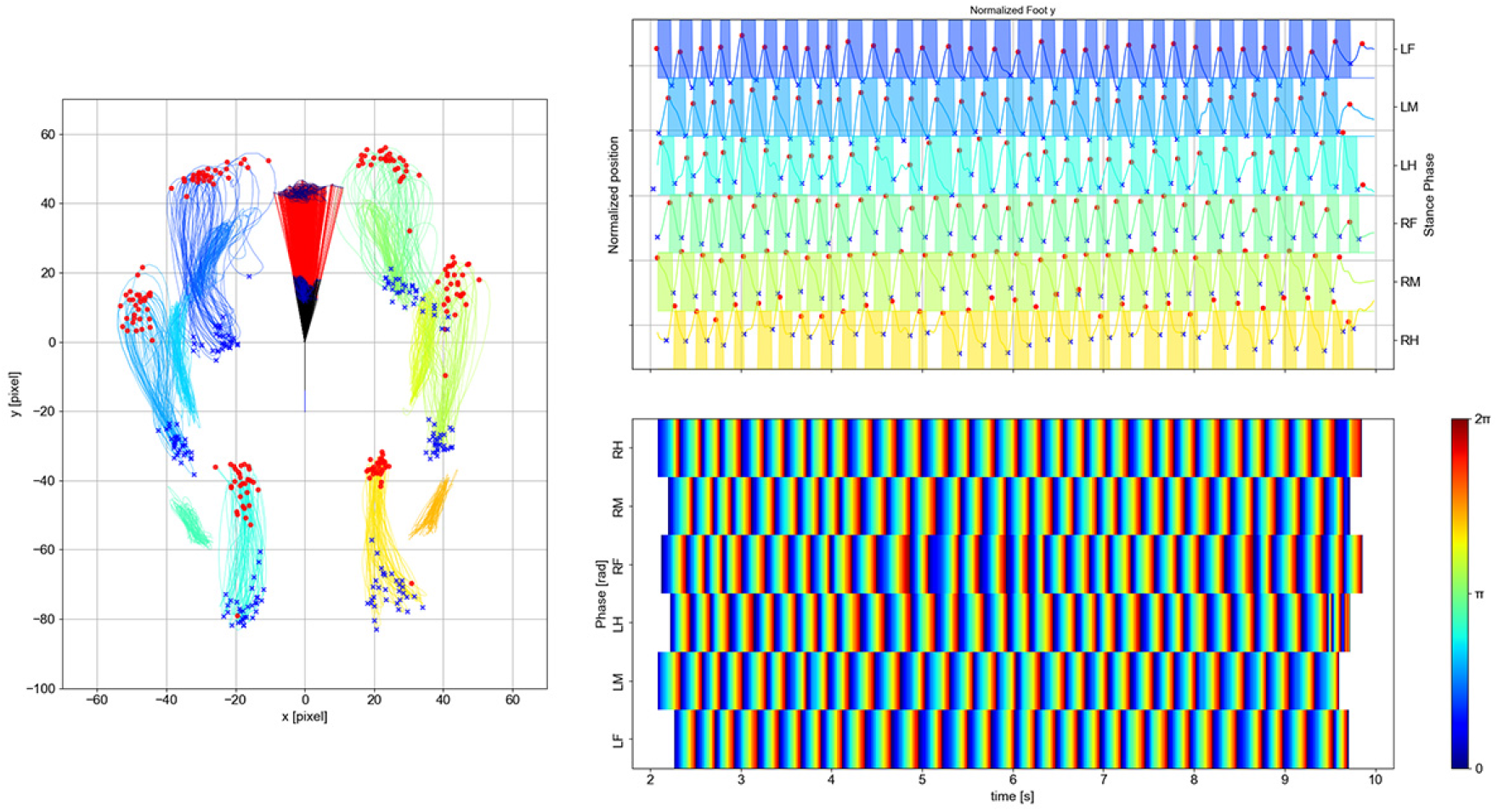
Extraction of Touchdown (TD) and Liftoff (LO) Events and Phase Estimation from Leg Trajectory Time Series. The upper-right panel shows an example of normalized time-series data of a leg tarsus, from which TD (red circles) and LO (blue crosses) events were extracted following the procedure described in “Materials and Methods, Gait Analysis” of the main text. Shaded regions indicate the stance phase, defined as the interval from TD to LO. The extracted TD and LO events are overlaid on the corresponding foot trajectories in the *x*–*y* plane of the right panel. For this trial, although some outliers in TD and LO detection appear near the termination of walking, such evident outliers were excluded from all statistical analyses. The lower-right panel shows the instantaneous phase of each leg, computed from foot trajectory data using the Hilbert transform and visualized as a colormap.

**Figure S5:**
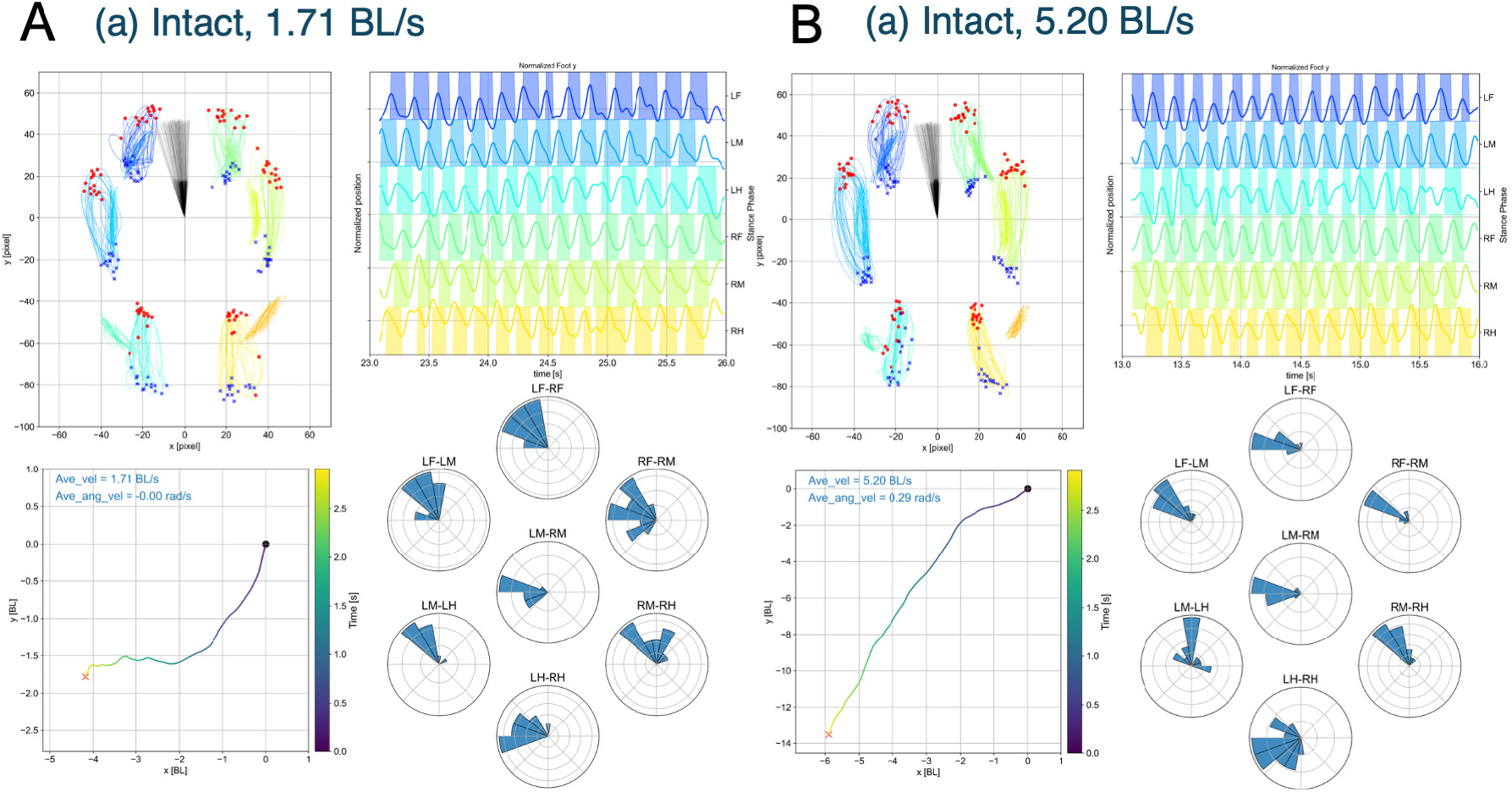
Representative Walking Patterns under the Intact Condition (a). Two trials with different mean forward velocities are shown: A, 1.71 BL s^−1^ (Movie SM2); B, 5.20 BL s^−1^ (Movie SM3). **Top left**: Trajectories of the Head, Pro, and the FTi–tarsus (e.g., LF2, LF1) positions of all legs plotted in the *x*–*y* plane (pixel units). The coordinate system is identical to that used in Figs. S3 and S4. Red circles indicate touchdown (TD) events, and blue crosses indicate lift-off (LO) events. **Top right**: Gait diagrams showing normalized y-coordinates of each leg, with stance phases (from TD to LO) highlighted. **Bottom left**: Reconstructed walking trajectories in the *x*–*y* plane. Black circles mark the starting positions, red crosses indicate the final positions, and the color scale represents elapsed time from start to end. Text annotations indicate the mean forward and rotational velocities for each trial. **Bottom right**: Inter-leg phase relationships at the TD timing of the LH leg, with phases extracted using the method described in Fig. S4.

**Figure S6:**
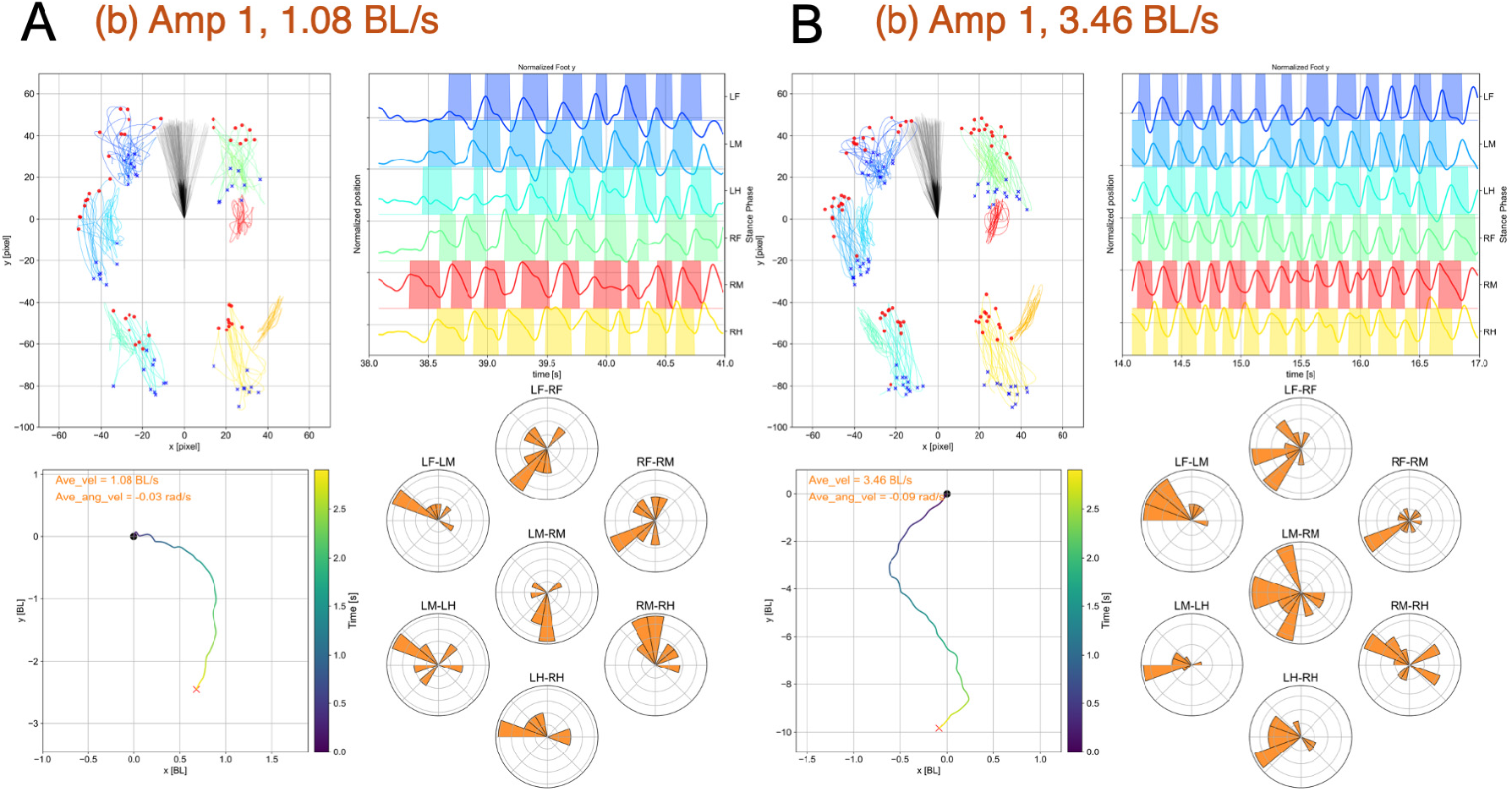
Representative Walking Patterns under the Amp 1 Condition (b). A, 1.08 BL s^−1^ (Movie SM4); B, 3.46 BL s^−1^ (Movie SM5).

**Figure S7:**
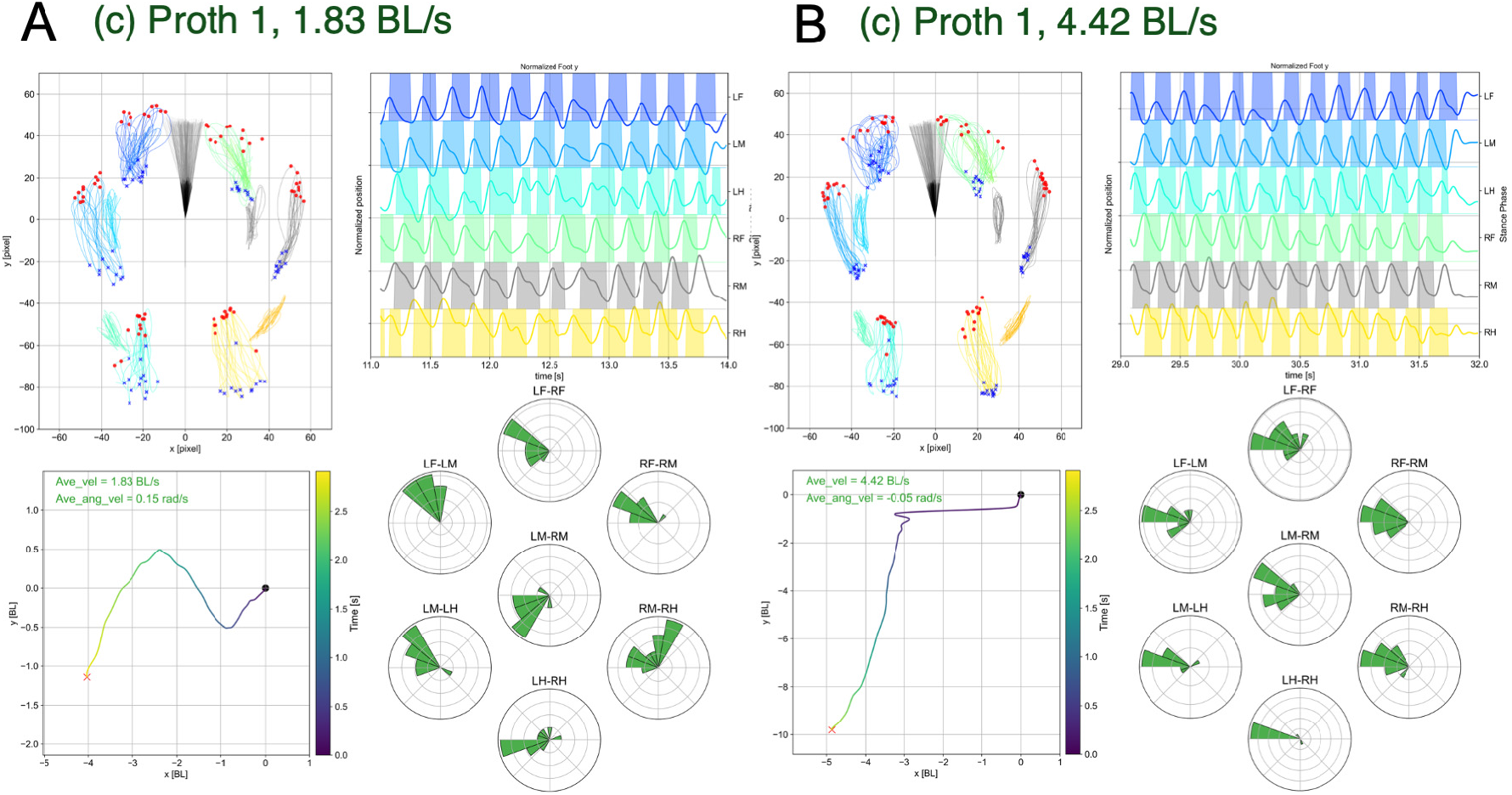
Representative Walking Patterns under the Proth 1 Condition (c). A, 1.83 BL s^−1^ (Movie SM6); B, 4.42 BL s^−1^ (Movie SM7).

**Figure S8:**
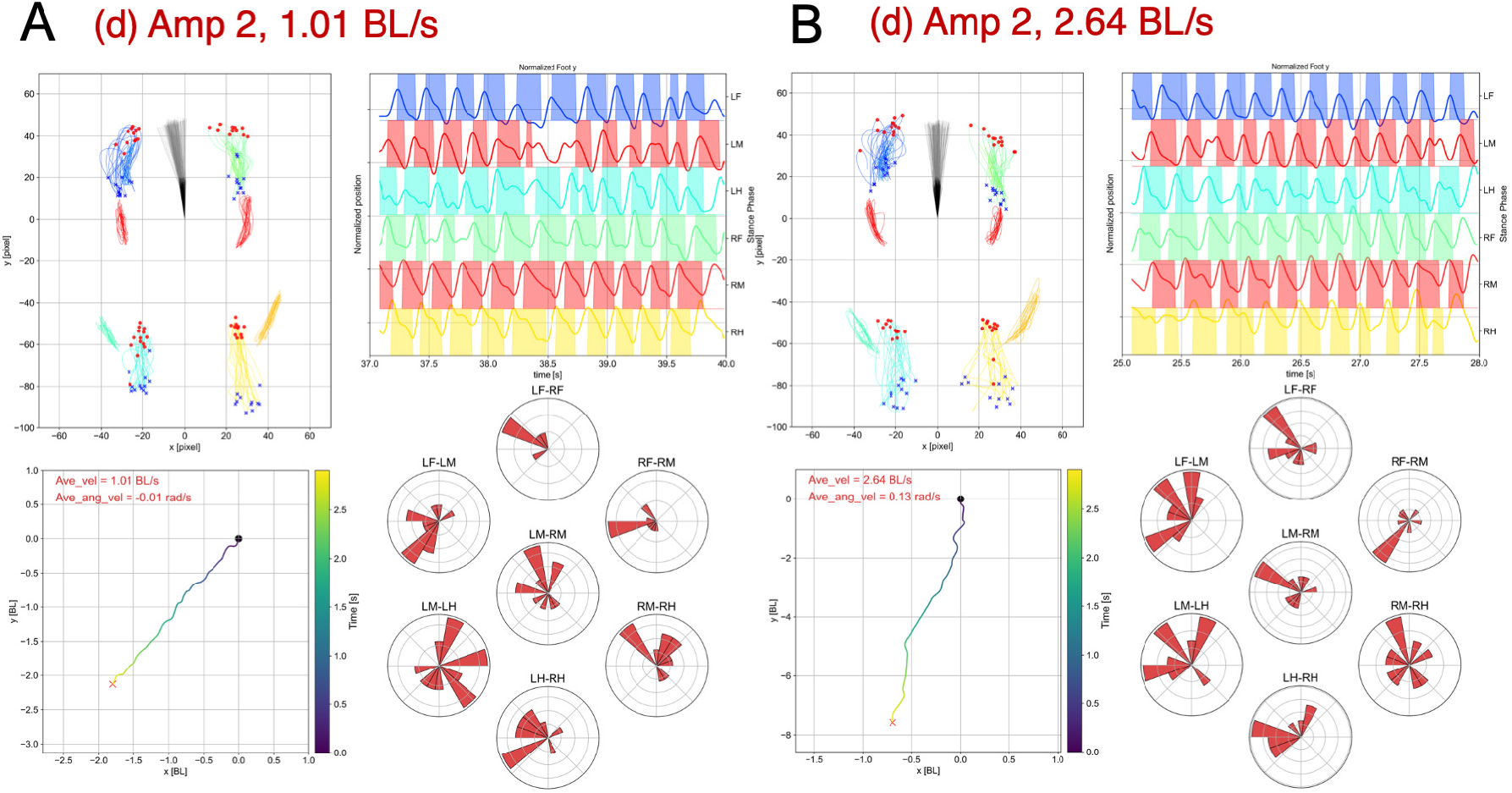
Representative Walking Patterns under the Amp 2 Condition (d). A, 1.01 BL s^−1^ (Movie SM8); B, 2.64 BL s^−1^ (Movie SM9).

**Figure S9:**
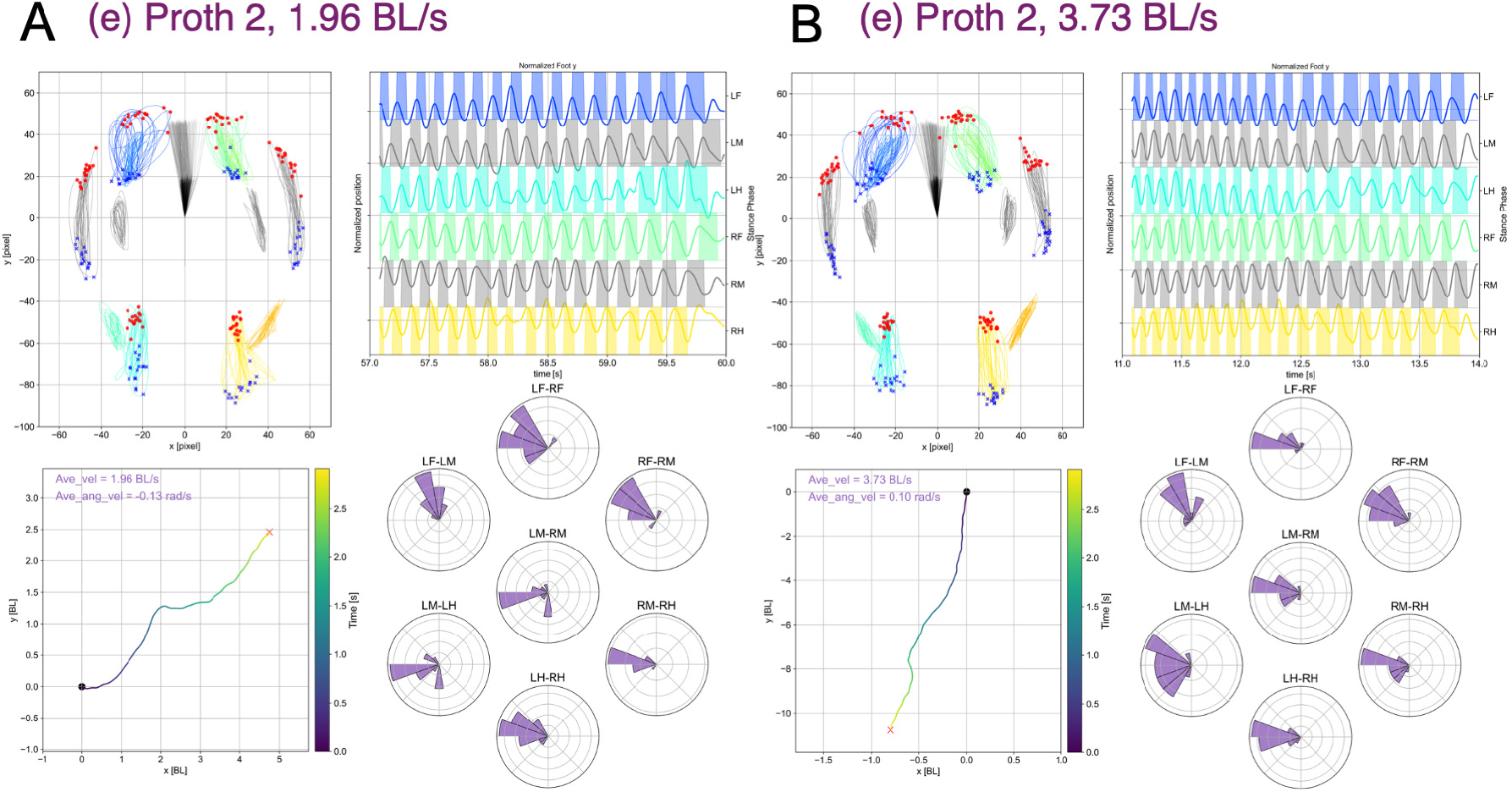
Representative Walking Patterns under the Proth 2 Condition (e). A, 1.96 BL s^−1^ (Movie SM10); B, 3.73 BL s^−1^ (Movie SM11).

**Figure S10:**
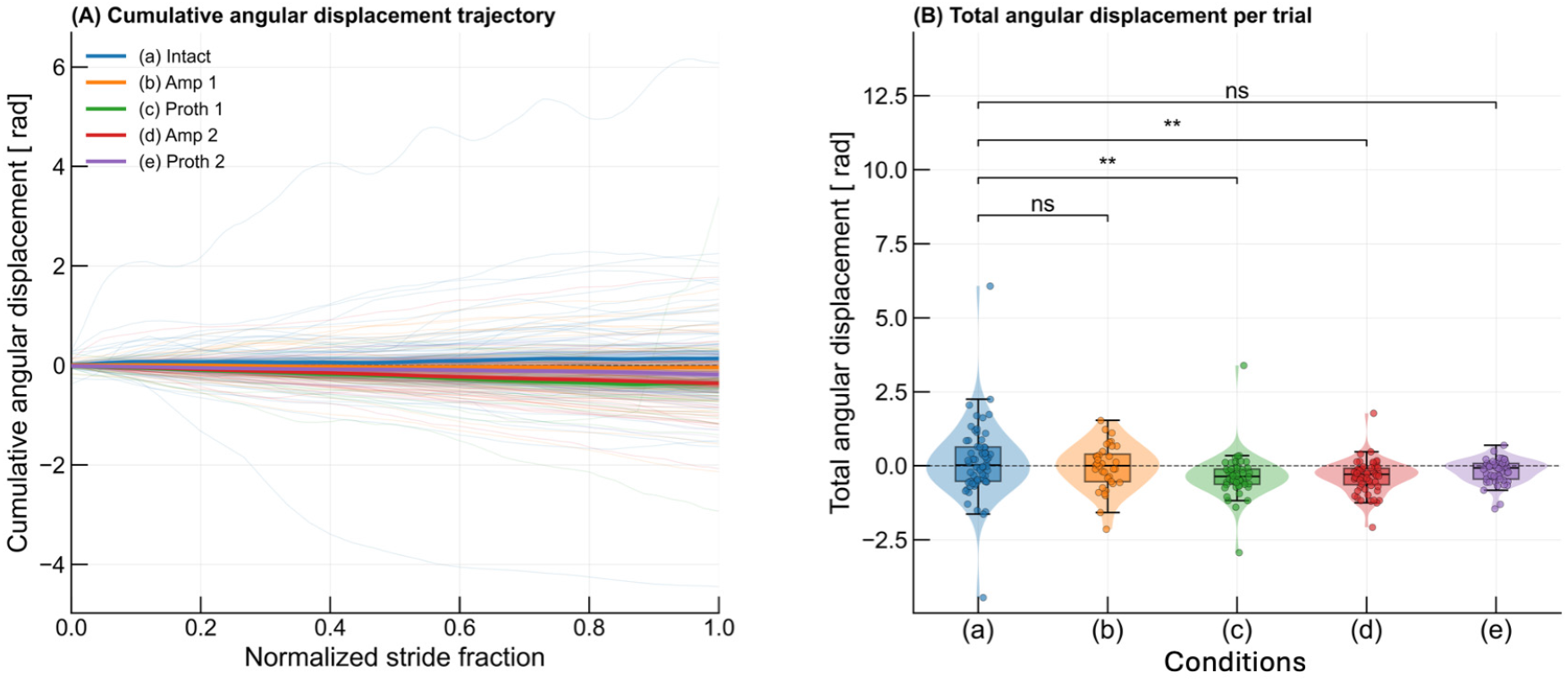
Cumulative angular displacement across five experimental conditions. (A) Trajectories for individual trials (thin lines) and condition means ± SEM (thick lines with shading), plotted against normalized stride fraction (0 = trial start; 1 = trial end). (B) Total cumulative angular displacement per trial (violin with embedded box plot and individual data points); the dashed line at zero indicates symmetric heading. BH-FDR-corrected Mann–Whitney U tests vs. the Intact condition: ^∗∗^, *p <* 0.01; ns, not significant. *n* = 64, 36, 49, 59, 49 trials for conditions (a)–(e), respectively.

**Table S1:** Watson–Williams tests (BH-FDR corrected across 42 tests) for inter-leg phase relationships in Fig. 2. Three representative leg pairs involving the prosthetic limb are shown; results for all seven pairs are indicated in Fig. 2.

| Leg pair | Comparison | Adj. $p$ | Sig. | DO |
| --- | --- | --- | --- | --- |
| *Intact (a) vs. manipulation conditions* |  |  |  |  |
| LM–RM | vs. (b) Amp 1 | $1.19 \times 10^{-12}$ | *** | 0.639 |
| LM–RM | vs. (c) Proth 1 | $8.48 \times 10^{-2}$ | ns | 0.877 |
| LM–RM | vs. (d) Amp 2 | $7.70 \times 10^{-43}$ | *** | 0.491 |
| LM–RM | vs. (e) Proth 2 | $1.40 \times 10^{-4}$ | *** | 0.657 |
| RF–RM | vs. (b) Amp 1 | $6.59 \times 10^{-19}$ | *** | 0.604 |
| RF–RM | vs. (c) Proth 1 | $1.95 \times 10^{-1}$ | ns | 0.865 |
| RF–RM | vs. (d) Amp 2 | $1.42 \times 10^{-5}$ | *** | 0.528 |
| RF–RM | vs. (e) Proth 2 | $7.03 \times 10^{-1}$ | ns | 0.694 |
| RM–RH | vs. (b) Amp 1 | $2.31 \times 10^{-3}$ | ** | 0.660 |
| RM–RH | vs. (c) Proth 1 | $5.71 \times 10^{-15}$ | *** | 0.701 |
| RM–RH | vs. (d) Amp 2 | $1.63 \times 10^{-3}$ | ** | 0.580 |
| RM–RH | vs. (e) Proth 2 | $7.01 \times 10^{-9}$ | *** | 0.652 |
| *Amputation vs. prosthesis (direct pairwise comparisons)* |  |  |  |  |
| LM–RM | (b) Amp 1 vs. (c) Proth 1 | $1.77 \times 10^{-10}$ | *** | — |
| LM–RM | (d) Amp 2 vs. (e) Proth 2 | $1.45 \times 10^{-13}$ | *** | — |
| RF–RM | (b) Amp 1 vs. (c) Proth 1 | $5.23 \times 10^{-10}$ | *** | — |
| RF–RM | (d) Amp 2 vs. (e) Proth 2 | $4.73 \times 10^{-3}$ | ** | — |
| RM–RH | (b) Amp 1 vs. (c) Proth 1 | $6.49 \times 10^{-2}$ | ns | — |
| RM–RH | (d) Amp 2 vs. (e) Proth 2 | $8.72 \times 10^{-5}$ | *** | — |

**Notes:** Watson–Williams two-sample tests compared mean inter-leg phase angles between conditions. All p-values are BH-FDR corrected across 42 pairwise tests (7 pairs × 6 comparisons). Significance codes: \*\*\*, $p < 0.001$ ; \*\*, $p < 0.01$ ; \*, $p < 0.05$ ; ns, not significant. DO: distribution overlap between Intact and the experimental condition (0–1); “—”: not applicable.

**Table S2:** Statistical comparisons for locomotor velocity and angular velocity in. **Fig. 3 (Mann–Whitney *U* test, two-sided, BH-FDR corrected across 12 tests).**

| Metric | Comparison | Condition | Raw $p$ | Adj. $p$ | Sig. |
| --- | --- | --- | --- | --- | --- |
| *Intact (a) vs. each condition* |  |  |  |  |  |
| Velocity | (a) vs. (b) | Amp 1 | $4.61 \times 10^{-3}$ | $1.11 \times 10^{-2}$ | * |
| Velocity | (a) vs. (c) | Proth 1 | $1.39 \times 10^{-4}$ | $5.57 \times 10^{-4}$ | *** |
| Velocity | (a) vs. (d) | Amp 2 | $< 1 \times 10^{-6}$ | $< 1 \times 10^{-6}$ | *** |
| Velocity | (a) vs. (e) | Proth 2 | $< 1 \times 10^{-6}$ | $< 1 \times 10^{-6}$ | *** |
| Angular velocity | (a) vs. (b) | Amp 1 | $6.54 \times 10^{-1}$ | $7.13 \times 10^{-1}$ | ns |
| Angular velocity | (a) vs. (c) | Proth 1 | $1.16 \times 10^{-3}$ | $3.49 \times 10^{-3}$ | ** |
| Angular velocity | (a) vs. (d) | Amp 2 | $1.21 \times 10^{-2}$ | $2.08 \times 10^{-2}$ | * |
| Angular velocity | (a) vs. (e) | Proth 2 | $2.88 \times 10^{-1}$ | $3.84 \times 10^{-1}$ | ns |
| *Amputation vs. prosthesis (direct pairwise comparisons)* |  |  |  |  |  |
| Velocity | (b) vs. (c) | Amp 1 vs. Proth 1 | $6.53 \times 10^{-1}$ | $7.13 \times 10^{-1}$ | ns |
| Velocity | (d) vs. (e) | Amp 2 vs. Proth 2 | $8.15 \times 10^{-1}$ | $8.15 \times 10^{-1}$ | ns |
| Angular velocity | (b) vs. (c) | Amp 1 vs. Proth 1 | $1.03 \times 10^{-2}$ | $2.06 \times 10^{-2}$ | * |
| Angular velocity | (d) vs. (e) | Amp 2 vs. Proth 2 | $2.72 \times 10^{-2}$ | $4.07 \times 10^{-2}$ | * |

**Notes:** Mann–Whitney $U$ tests (two-sided) applied to per-recording-session median values. All 12 p-values corrected jointly using BH-FDR. Significance codes: \*\*\*, $p < 0.001$ ; \*\*, $p < 0.01$ ; \*, $p < 0.05$ ; ns, not significant.

**Table S3:** Body length and weight of individual crickets used in the experiments.

| Animal ID | Body length (mm) | weight (g) |
| --- | --- | --- |
| cricket017 | 25.30 | 0.98 |
| cricket018 | 25.75 | 0.97 |
| cricket019 | 25.32 | 0.95 |
| cricket020 | 25.90 | 1.17 |
| cricket021 | 25.26 | 0.93 |

**Table S4:** Two-sample Kolmogorov–Smirnov (KS) test of foot-placement distributions (stride-level sampling), each condition vs. Intact, for the lateral (*x*) and anteroposterior (*y*) axis of each leg. *D* is the KS effect size (maximum difference between the cumulative distributions); DO is the distribution overlap with Intact; SH-DO is the Intact split-half reference (natural within-condition variability). Amputated legs are omitted per condition.

| Condition | Leg | Lateral $x$ | | | Anteroposterior $y$ | | |
| --- | --- | --- | --- | --- | --- | --- | --- |
| | | SH-DO | DO | $D$ | SH-DO | DO | $D$ |
| (b) Amp 1 | LF | 0.900 | 0.860 | 0.232 | 0.891 | 0.819 | 0.293 |
|  | LM | 0.953 | 0.934 | 0.066 | 0.947 | 0.850 | 0.523 |
|  | LH | 0.917 | 0.879 | 0.165 | 0.911 | 0.906 | 0.186 |
|  | RF | 0.851 | 0.792 | 0.072 | 0.935 | 0.841 | 0.413 |
|  | RH | 0.929 | 0.628 | 0.363 | 0.912 | 0.894 | 0.230 |
| (c) Proth 1 | LF | 0.900 | 0.884 | 0.062 | 0.891 | 0.795 | 0.354 |
|  | LM | 0.953 | 0.945 | 0.071 | 0.947 | 0.821 | 0.507 |
|  | LH | 0.917 | 0.823 | 0.188 | 0.911 | 0.895 | 0.259 |
|  | RF | 0.851 | 0.863 | 0.080 | 0.935 | 0.779 | 0.542 |
|  | RM | 0.883 | **0.319** | **0.600** | 0.938 | 0.916 | 0.194 |
|  | RH | 0.929 | 0.726 | 0.199 | 0.912 | 0.895 | 0.235 |
| (d) Amp 2 | LF | 0.900 | 0.724 | 0.214 | 0.891 | 0.777 | 0.499 |
|  | LH | 0.917 | 0.777 | 0.213 | 0.911 | 0.764 | 0.459 |
|  | RF | 0.851 | 0.704 | 0.287 | 0.935 | 0.761 | 0.558 |
|  | RH | 0.929 | 0.714 | 0.397 | 0.912 | 0.818 | 0.403 |
| (e) Proth 2 | LF | 0.900 | 0.801 | 0.197 | 0.891 | 0.792 | 0.459 |
|  | LM | 0.953 | **0.246** | **0.593** | 0.947 | 0.899 | 0.127 |
|  | LH | 0.917 | 0.644 | 0.455 | 0.911 | 0.834 | 0.388 |
|  | RF | 0.851 | 0.899 | 0.175 | 0.935 | 0.822 | 0.467 |
|  | RM | 0.883 | **0.253** | **0.684** | 0.938 | 0.902 | 0.163 |
|  | RH | 0.929 | **0.493** | **0.587** | 0.912 | 0.884 | 0.343 |

**Notes:** Stride-level sample sizes were large ( $n \approx 1,300$ –2,600 stride placements per leg and condition); accordingly *all* 42 KS comparisons were significant after BH-FDR correction ( $q < 0.001$ , \*\*\*), so significance is not tabulated separately. Recovery is therefore read from the effect size $D$ and from DO relative to the leg-specific SH-DO reference: small $D$ with DO near SH-DO indicates preservation, whereas large $D$ with DO well below SH-DO indicates a genuine deficit exceeding natural within-condition variability. Bold entries mark the pronounced lateral ( $x$ ) foot-placement deficits of the prosthetic middle legs (RM, LM) and load-bearing hindlegs; note that the anteroposterior ( $y$ ) placement of the same prosthetic legs remains close to the SH-DO reference.

